# PURELIGHT: a quantitative photon-counting framework unifying intensity and lifetime imaging at video rate across detector technologies

**DOI:** 10.64898/2026.08.28.747260

**Authors:** Felipe Velasquez Moros, Dorian Amiet, Daria Skwarzynska, Rachel Meister, Urvashi Dalvi, Saidong Ma, Peter Rupprecht, Shuting Han, Philipp Bethge, Lukas Krainer, Giulia Acconcia, Ivan Rech, Fritjof Helmchen, Paul Zbinden, Aiman S. Saab, Bruno Weber, Luca Ravotto

## Abstract

Quantitative fluorescence microscopy requires photon-efficiency, speed and accurate intensity and lifetime measurements. Time-correlated single-photon counting (TCSPC) simultaneously captures intensity and lifetime, but photon pile-up distorts both signals at high count rates, preventing fast acquisitions. Existing corrections discard photons, distort intensity, or require specialized detectors. Here we introduce PURELIGHT, an integrated hardware and software framework that simultaneously recovers undistorted intensities and lifetimes at count rates far beyond conventional pile-up limits. PURELIGHT works with hybrid photodetectors, silicon photomultipliers and photomultiplier tubes while retaining over three times more photons than alternative approaches. Using two-photon imaging, we showcase PURELIGHT’s superior accuracy and spatial contrast, demonstrating video-rate subcellular lifetime imaging in awake mice, a unique lifetime-calibrated ratiometric modality and crosstalk-free temporal multiplexing. By removing the limits that have confined TCSPC to low-signal applications, PURELIGHT promotes the adoption of quantitative, photon-efficient microscopy across the life sciences.

## 1 Introduction

Following its invention over three decades ago [1], two-photon fluorescence microscopy (2PM) has become central to neuroscience, enabling *in vivo* measurements of neuronal and glial activity, metabolism, blood flow and neurotransmission [2]. Despite these successes, accurate quantitative imaging of the biochemical processes driving brain function is still in its infancy. The main challenge lies in the photodetection chain, which should simultaneously deliver photon efficiency, acquisition speed and accurate readouts of both fluorescence intensity and lifetime. No established technology provides all three.

Most 2PM experiments rely on the temporal integration of a voltage signal from photomultiplier tubes (PMTs), a simple strategy that tolerates high photon fluxes but adds electronic noise and does not provide photon timing information. Both limitations are becoming more consequential. On one hand, as the field evolves towards ever faster acquisitions, a very small number of photons is collected in each pixel [3, 4]; in this regime, any noise added by the detection chain directly limits measurement accuracy. On the other hand, the lack of timing information prevents the implementation of Fluorescence Lifetime Imaging (FLIM), the state-of-the-art method to extract quantitative biological information from thick tissues and living animals. In fact, while intensity is subject to systematic distortions due to factors like scattering, absorption and uneven sensor expression, FLIM is free from these confounds and extracts significantly more information from the same fluorescence signal [5, 6].

Time-correlated single photon counting (TCSPC) can overcome the noise and quantitation limits of analog integration. By detecting and timing each individual photon, TCSPC provides shot-noise-limited intensity detection [7, 8] together with intensity-independent FLIM readouts [9], and integrates seamlessly with the pulsed excitation sources already used in 2PM. Its accuracy, however, is bounded by the so-called “pile-up” effect: after a photon is detected, the system is unable to collect new photons for some time, compromising both intensity and lifetime accuracy [10]. Even with modern electronics and state-of-the-art detectors, these distortions become significant once the incident photon rate exceeds 5–10% of the excitation laser repetition rate. In a typical biological image, the need to avoid pile-up in the brightest areas forces a reduction in signal across the entire image, so that acquiring a noise-robust FLIM image requires minutes and precludes the study of fast biological processes.

In recent years, several approaches have increased the usable count rates of TCSPC systems. Photon-rejection and imposed-blind-time strategies [11, 12] can preserve fluorescence decays but discard a substantial fraction of photons and often do not recover intensity quantitatively [11]. Model-based corrections [13] can account for pile-up during fitting but increase computational complexity and constrain downstream analysis, whereas specialized detector architectures [14] may offer high performance but are not readily transferable to the large-area detectors commonly used in scattering tissue. Thus, a general strategy that simultaneously preserves photon efficiency, intensity and lifetime accuracy, and acquisition speed across detector technologies has remained unavailable.

Here we introduce PURELIGHT (Pile-Up Removal for Enhanced Lifetime and Intensity Gathering in High-flux TCSPC), an acquisition and correction framework that combines fast digitization with an extended time-tagged data format recording pulse shape information for each photon detection. These measurements allow PURELIGHT to recover quantitative intensity and undistorted fluorescence lifetime while retaining substantially more photons than rejection-based approaches. We validate PURELIGHT across detectors with different response speed and noise characteristics, such as hybrid photodetectors (HPDs), silicon photomultipliers (SiPMs) and PMTs, demonstrating quantitative subcellular imaging in biological tissue and awake mice, including video-rate calcium imaging, combined ratiometric and lifetime measurements and low-crosstalk temporal multiplexing. By extending accurate photon counting across the wide range of signal levels encountered in biological microscopy, PURELIGHT transforms TCSPC from a specialized low-flux FLIM method into a practical and broadly applicable detection strategy for high-speed, low-noise quantitative microscopy.

## 2 Results

### 2.1 PURELIGHT enables accurate FLIM and intensity corrections at high count rates *in silico*

In TCSPC, the arrival time of a photon is typically determined as the time at which a signal from a photodetector crosses a predetermined voltage threshold. Only once the signal returns below the threshold, the detection of a new photon is possible. At high count rates, one or more photons can hit the detector while the signal is still above threshold (“pile-up”, Figure 1a), causing the loss of detection events and temporal distortions in their arrival time profile.

**Fig. 1.**
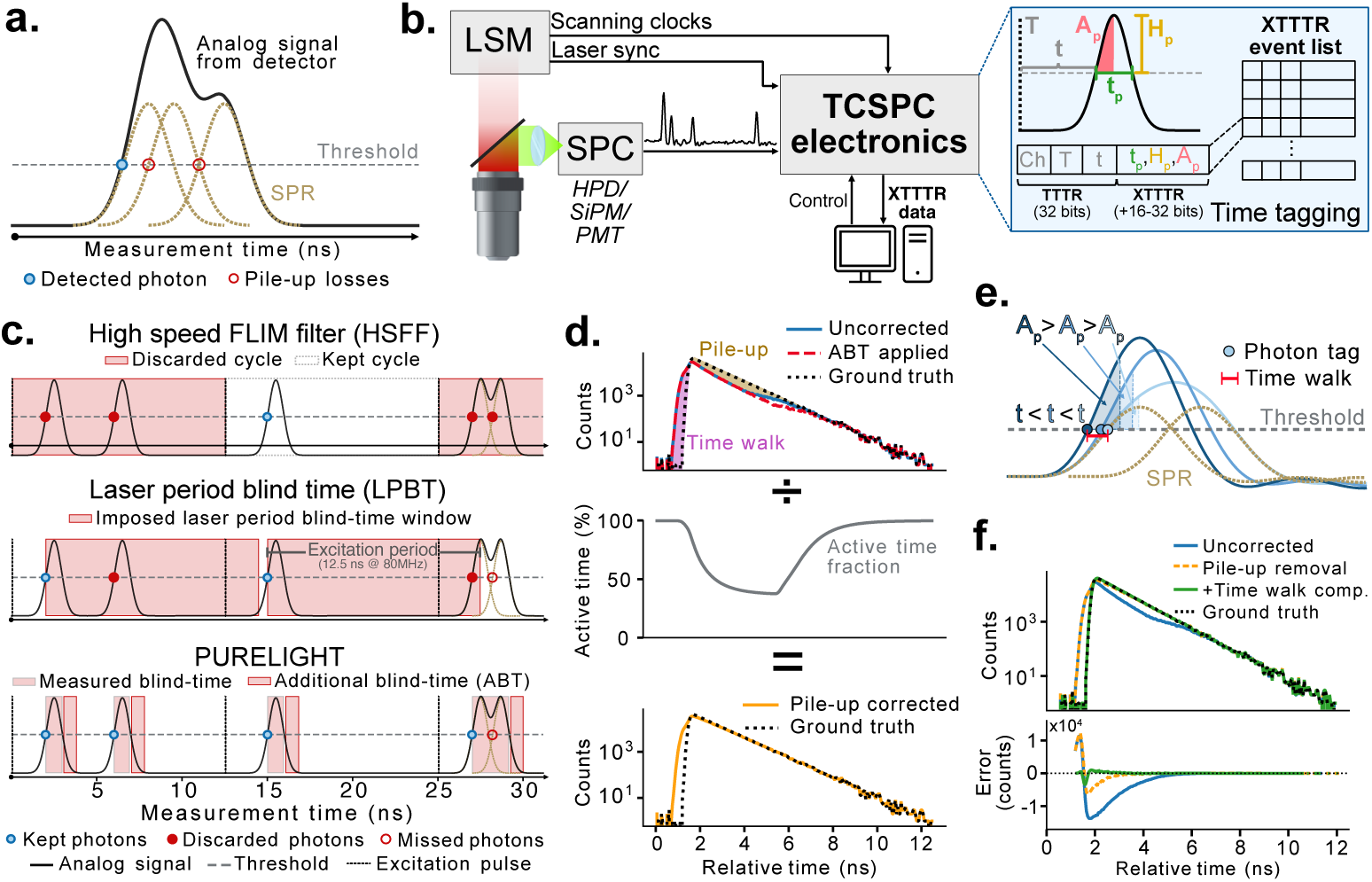
TCSPC distortions, data acquisition, and processing. **a)** Pile-up losses in threshold based TCSPC systems. Multiple photons reaching the detector in a short period of time combine in the analog output of the detector (black) and result in a single threshold crossing. **b)** XTTTR data collection from a laser scanning microscope. Signals from up to 7 single photon counting detectors, laser sync, and microscope scanning clocks are digitized and processed on a field-programmable gate array (FPGA) to extract the XTTTR data: absolute time (*T*: excitation cycle number), relative time (*t*: time delay after the excitation pulse), maximum value above the threshold (*H_p_*), time above the threshold (*t_p_*), and area under the rising edge (*A_p_*). The collected XTTTR data is transmitted to the PC for further analysis and concurrently processed on the FPGA to provide real-time intensity and pseudo-FLIM images for all active channels. **c)** Photon detection with different pile-up correction strategies. The high-speed FLIM filter (HSFF, top) [11] only records events from excitation cycles with exactly one detected photon. The laser period blind time method (LPBT, middle) [12] discards all events after a detected event for a duration equal to the excitation period. PURELIGHT (bottom) keeps all detected events and measures the time above threshold. Then, a fixed added blind time (ABT) is imposed to eliminate falling edge effects. **d)** Pile-up correction based on the active time fraction. Each time point in the pile-up distorted decay with ABT applied (red, top) gets divided by the percentage of time the system was able to detect photons at that time point (middle), resulting in a decay (orange, bottom) without the distortions generated by pile-up losses. **e)** Origin of time walk in fixed threshold systems. When multiple photons are detected in rapid succession, i.e. closer than the single photon response (SPR) rise time, the analog output of the detector combines their individual contributions, resulting in a faster rising signal and an earlier threshold crossing. The shaded area corresponds to the measured *A_p_* for each example detection. **f)** Complete PURELIGHT correction on a simulated decay (Gaussian SPR 2 ns FWHM, *τ* = 1 ns, 80 Mcps). Combining the pile-up loss corrected curve (orange) with a time walk compensation algorithm (Supplementary Note 1) results in a corrected decay (green) that closely matches the ground truth.

To recover the undistorted measurement, we introduce a novel time-tagging strategy that uses custom FPGA-based electronics equipped with fast ADCs to digitize the voltage pulses from photon counting detectors (Figure 1b, Supplementary Figure 1). This method, named eXtended TTTR (XTTTR), combines threshold crossing-based time tagging [15] with additional information such as the peak voltage (*H_p_*), the time spent above the detection threshold (*t_p_*) or the area of the signal for a given time after threshold crossing (*A_p_*).

The additional information contained in the XTTTR format allows us to implement multiple post-processing pile-up correction algorithms, such as the HSFF [11] available in commercial microscopes or our previously reported LPBT method [12]. However, while these methods can correct lifetime distortions, they discard a large fraction of collected photons (Figure 1c), and either cannot correct intensity or require expensive state-of-the-art Hybrid Photodetectors (HPDs).

To overcome these limitations, we introduce PURELIGHT, a comprehensive and detector-agnostic method to correct pile-up losses and timing distortions while discarding far fewer photons than alternative methods (Figure 1d-f).

The correction algorithm consists of three steps (Supplementary Note 1). First, to remove distortions originating from the interaction between falling and rising edges of two detection events, the blind time (i.e. the time over threshold *t_p_*, in which the detection of an incoming photon is not possible) of each event is elongated by a predetermined small amount and any detected events within this extended period are removed (Figure 1c). Then, the detection efficiency of the system at each time delay relative to the excitation pulse (relative time, *t*) is computed and used as a scaling factor to retrieve the real signal intensity (Figure 1d) [16]. Finally, to compensate for the time walk induced by overlapping pulses (Figure 1e,f), the area under the rising edge of the event (*A_p_*) is used to compute a time shift based on a calibration measurement, mimicking a constant fraction discriminator (CFD), but operating on a metric more robust to partially overlapping events.

We first validated PURELIGHT by performing Monte-Carlo simulations of pileup affected photon arrival times based on monoexponential fluorescence decays under different scenarios in terms of count rates and detector pulse characteristics (Figure 2a top, Supplementary Figure 2).

**Fig. 2.**
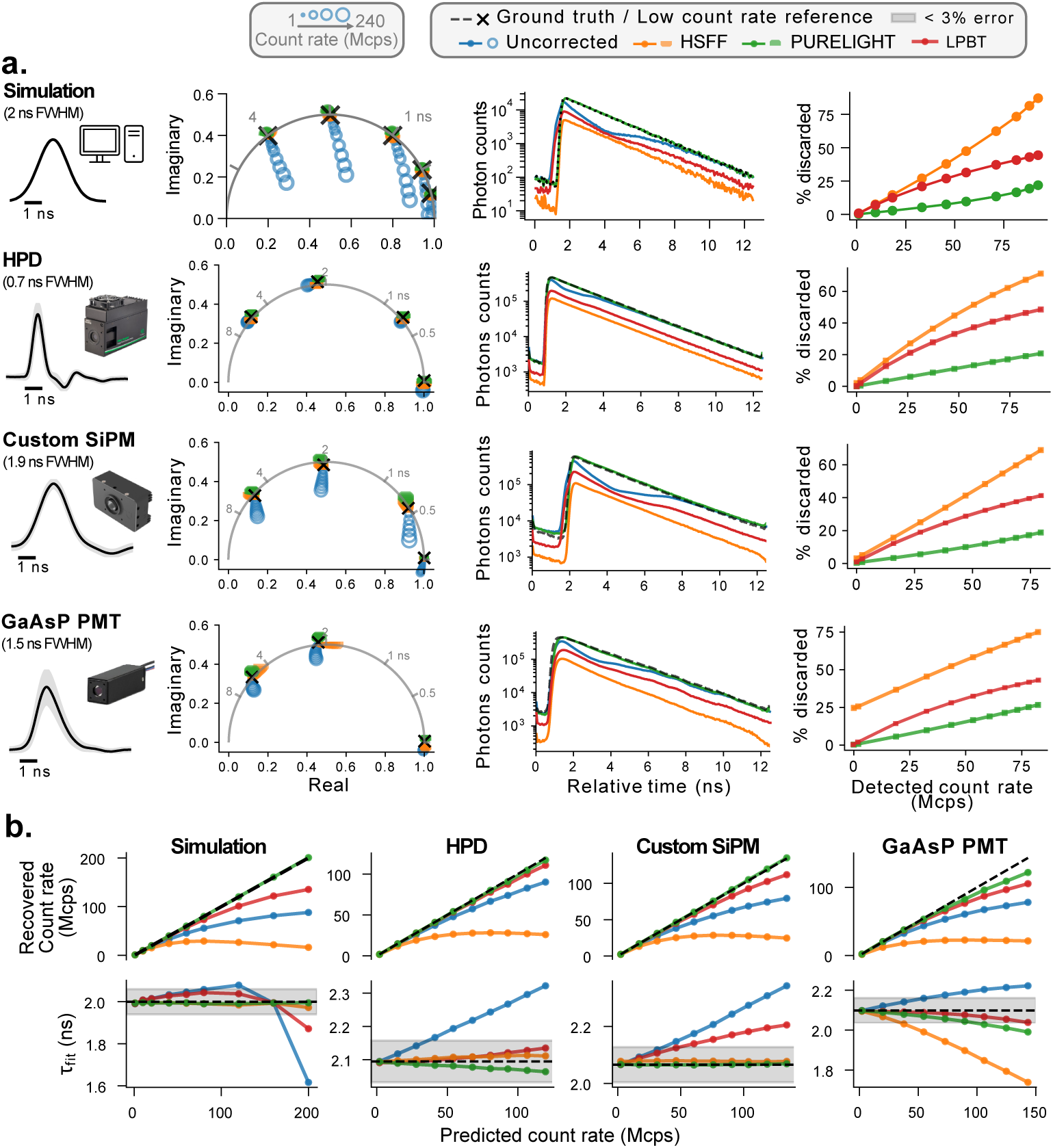
XTTTR data enables accurate FLIM and intensity measurements at high count rates. **a)** Validation of the HSFF, LPBT, and PURELIGHT corrections applied to simulated (detector response: Gaussian 2 ns FWHM) and experimental XTTTR data collected from multiple detector technologies using mono-exponential samples with various lifetimes. Rows, from top to bottom: simulation, HPD, custom SiPM and GaAsP PMT. The black traces (left) show each detector’s single photon response with the shaded area denoting its standard deviation. The phasor plots (2nd column) compare the stability of the HSFF (orange) and PURELIGHT (green) corrections to a low count rate reference (black) as the incident count rate increases (increasing marker size). The FLIM decays (3rd column) correspond to the XTTTR data from the *∼*2 ns sample at low count rate (dashed black line) and over 100 Mcps (colored lines) processed with the same methods as above. The counting efficiency plots (4th column) show the percentage of detected events that are discarded by each correction method as a function of count rate. **b)** Recovered count rate (intensity) and fitted lifetime dependence on count rate before and after correction for the same quenched fluorescein solution as above (additional lifetimes in Supplementary Figures 3-5) at up to 200% excitation rate (160 Mcps). Columns, from left to right: simulation, HPD, custom SiPM and GaAsP PMT.

Figure 2a (top row) shows a comparison of the PURELIGHT, HSFF, and LPBT corrections on simulated TCSPC data with a Gaussian pulse shape with 2 ns full width at half-maximum (FWHM). Both PURELIGHT and HSFF can reconstruct the temporal profiles accurately and achieve virtually undistorted phasor plots at up to about 300% of the excitation rate. As has been shown before, the LPBT correction requires a fast detector response in order to remove pile-up effectively [12] and thus it was not able to recover the undistorted temporal decay and accurate lifetime estimation in this simulation (Figure 2a, top row; Figure 2b, left). While the HSFF provides undistorted decays, PURELIGHT does so with a much greater photon efficiency: when the detection rate exceeds 100% (80 Mcps) the HSFF correction discards over 75% of the detected events while PURELIGHT only discards around 20%. In other words, with identical acquisition times, PURELIGHT can collect over three times the number of photons as HSFF.

### 2.2 PURELIGHT provides undistorted TCSPC across multiple detector technologies

To assess the real-world effectiveness of PURELIGHT in correcting pile-up distortions in a detector-agnostic way, we performed *in vitro* validations using three different detector technologies: a commercial HPD, a custom 3*×*3 mm^2^ Silicon Photomultiplier (SiPM, Methods), and a commercial GaAsP PMT. For each detector, we acquired TCSPC images of dye solutions with different lifetimes coupling our custom-made electronics to a two-photon microscope equipped with an 80 MHz femtosecond-pulsed laser. To test the accuracy and limits of pile-up correction algorithms, we excited the samples with progressively higher laser powers, with acquisitions at count rates below the pile-up limit used as ground truth reference. To demonstrate the robustness of the results independently of the chosen evaluation metric, we analyzed the results using visual inspection of the decays and residuals, phasor analysis, exponential fitting, and the method of moments (MoM) (Figure 2a; Supplementary Figures 3-5).

For all detectors, the phasor and decay plots (Figure 2a) show close alignment between the low and high count rate measurements after applying the PURELIGHT correction. Conversely, the HSFF method fails for PMTs, likely because it relies on the accurate classification of single-photon events, which is compromised by the larger pulse-to-pulse variability of PMTs. The LPBT correction works well for HPDs and PMTs (0.7 and 1.5 ns pulse FWHM respectively), while for the SiPM (1.9 ns pulse FWHM) there is a visible distortion in the decay.

Residuals inspection (Supplementary Figures 3-5) and exponential fitting (Figure 2b) show that PURELIGHT maintains the expected lifetime accuracy across a wide range of count rates for all detectors. The HSFF method performs well in cases where single photon events are clearly identifiable (simulation, HPDs, and SiPMs) while the LPBT method only provides accurate results when the detector response is fast (HPDs, PMTs).

For HPDs, the fitted lifetime drift with PURELIGHT remained below 60 ps across all tested lifetimes up to 200% of the excitation rate (*>*160 Mcps), and the MoM showed a drift of less than 17 ps. Our custom SiPM, despite its slower temporal response and higher dark counts, handled the highest count rates of all tested detectors (up to 280 Mcps, 350% of the excitation rate) while maintaining less than 50 ps of fitted lifetime drift and accurate photon counting linearity, offering the best dynamic range among the three detectors. PMTs, although more limited in FLIM accuracy due to their broader and more variable pulse shape, still benefited from the PURELIGHT and LPBT corrections: fitted lifetime drift remained below 3% up to 100% of the excitation rate (*∼*80 Mcps). For applications where peak count rates do not exceed 1 photon per pulse (10*×* the classic pile-up limit), PMTs offer a practical and cost-effective alternative to HPDs, particularly given their prevalence in existing microscopy systems and their low dark counts.

In comparison to other methods, PURELIGHT corrects lifetime distortions while at the same time providing correct intensity readouts (Figure 2b) and discarding significantly fewer photons (Figure 2a, right column). Even when compared with LPBT using HPD detectors (the most favorable scenario for LPBT), PURELIGHT achieves the same correction capability while discarding fewer than half as many photons.

### 2.3 PURELIGHT improves quantitative imaging at all light intensity regimes using low-cost detector technologies

The compatibility of PURELIGHT with SiPM and PMT detectors is a crucial advantage for TCSPC adoption, as it enables the construction of cost-effective and compact multichannel instruments. To demonstrate the practical utility of a fully corrected multichannel TCSPC microscope in a biologically relevant context, we performed two-photon FLIM imaging of two acute *ex vivo* preparations (Figure 3a).

**Fig. 3.**
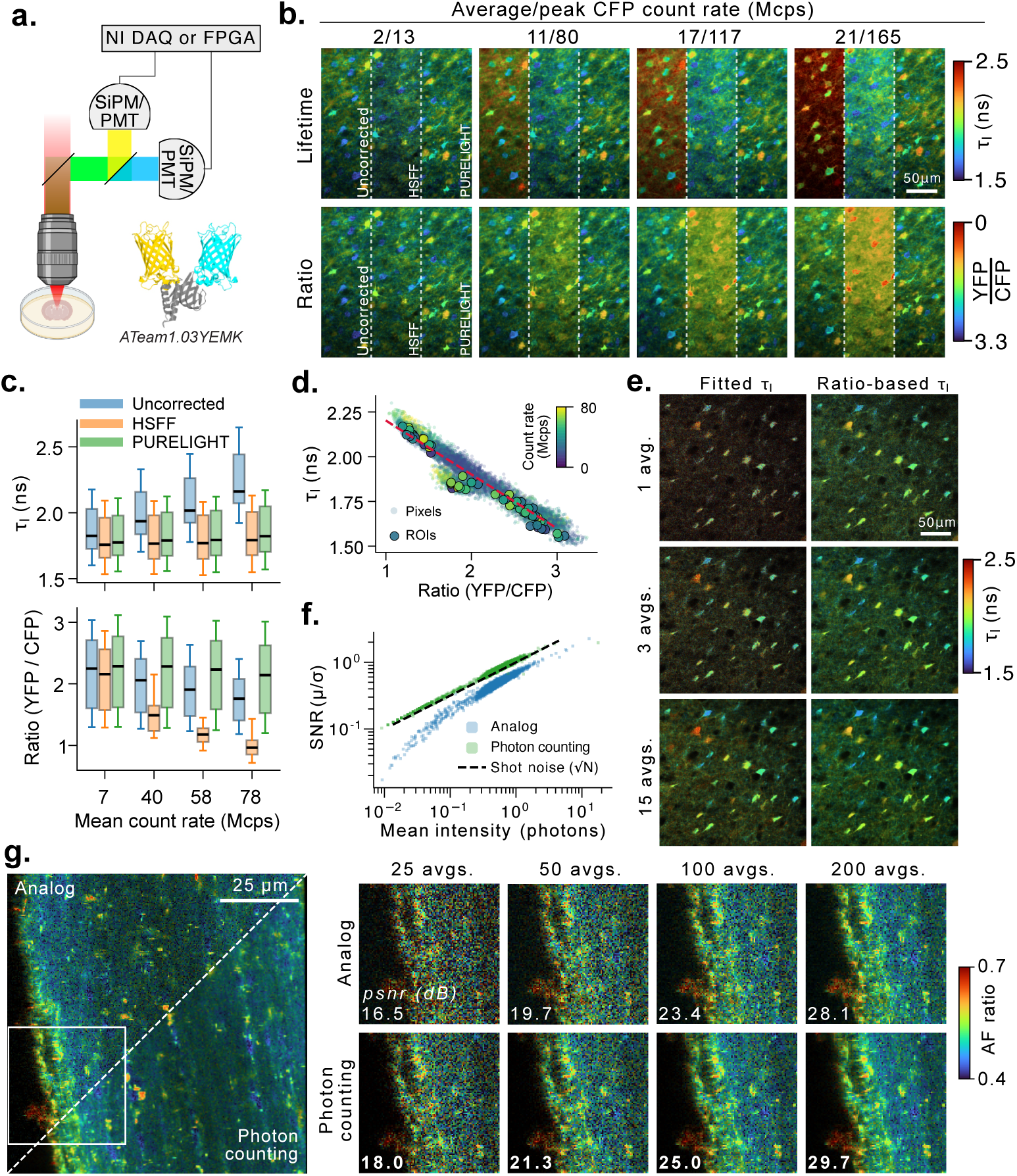
Simultaneous FLIM & ratiometric imaging with dual channel TCSPC. **a)** 2PM imaging of the neuron-targeted ATP sensor ATeam1.03YEMK in acute mouse cortical slices. Two SiPM modules or PMTs were used to detect cyan fluorescent protein (CFP – 475/50 nm) and yellow fluorescent protein (YFP – 542/50 nm) fluorescence in parallel. The output of each detector was either sent to the custom TCSPC electronics (photon counting) or the NI DAQ (analog). **b)** Comparison of pile-up correction methods on the intensity-weighted mean lifetime *τ_I_* (top) and ratiometric (bottom) images at different count rates (achieved by varying excitation power). **c)** ROI lifetime (top) and ratio (bottom) distributions at different count rates. **d)** Experimental correlation between YFP/CFP ratio and CFP lifetime. A linear fit of pixel values with varying ratios/lifetimes (red) provides a way to convert ratiometric images to lifetime. **e)** Lifetime images obtained from global fitting (left) and using the YFP/CFP ratio and lifetime correlation from panel d (right) using different number of averaged frames. At low number of averages (1 & 3 frames averaged) global fitting is unable to estimate lifetimes within the expected range (1.5 ns to 2.5 ns) for most pixels in the image. **f)** Signal-to-noise ratio (SNR) as a function of pixel intensity for analog integration (blue) and photon counting (purple) images. **g)** Autofluorescence imaging of a mouse optic nerve. The same acquisition was performed both with PMT detectors via analog integration (top) and photon counting (bottom). Averaging 400 frames (left) provides a ground truth image to evaluate the quality of fewer averaged frames (close-ups, right).

First, we imaged acute brain slices expressing the ATP sensor ATeam1.03 YEMK [17] using two of our custom SiPMs. By varying the excitation power we acquired images at progressively increasing count rates and compared the average CFP lifetime and YFP/CFP ratio (Figure 3b). Both PURELIGHT and HSFF maintained stable lifetime estimates across the full range of count rates, while the uncorrected data showed a progressive drift. However, the HSFF correction distorted the intensity of each channel, causing the YFP/CFP ratio to drift with increasing count rate despite the correct lifetime values. Only the PURELIGHT correction preserved both the life-time and the ratiometric readout across the full range of count rates, enabling fast and accurate imaging of ATP dynamics (Figure 3c).

The simultaneous correction of intensity and lifetime offered by PURELIGHT allows us to exploit the relationship between YFP/CFP ratio and intensity-weighted lifetime to combine the quantitative accuracy of FLIM with the better photon economy of ratiometric imaging. In fact, it can be demonstrated that the apparent *K_d_* of the sensor is the same between the two modalities [18], and thus there is a linear relationship between the two quantities (Figure 3d), that can be leveraged to generate lifetime images with a much better signal-to-noise ratio than what is obtainable from direct fitting (Figure 3e).

To demonstrate the superiority of a TCSPC microscope with respect to its analog counterpart under low light intensity regimes we performed ratiometric autofluorescence imaging using PMT detectors. We imaged an optic nerve preparation under 800 nm two-photon excitation, mimicking a typical low-light scenario encountered in label-free metabolic imaging. As expected by the shot-noise limited nature of the TCSPC signal (Figure 3f), for a similar number of photons reaching the detector, TCSPC produced images with higher peak signal-to-noise ratio, facilitating the visualization of finer morphological features (Figure 3g).

### 2.4 PURELIGHT enables video-rate subcellular imaging of astrocytic and neuronal calcium concentrations in awake mice

Astrocytic calcium activity plays a central role in neurovascular coupling, synaptic modulation, and the response to pathological events such as epileptic seizures [19–22]. Calcium dynamics span a wide range of spatial scales, ranging from subcellular microdomains in fine processes to large waves propagating across brain areas, and evolve on a sub-second to second timescale [23, 24]. Capturing these dynamics quantitatively requires imaging at high detection count rates, which inevitably leads to pile-up.

To demonstrate the capabilities of PURELIGHT in such a demanding biological scenario, we performed two-photon FLIM imaging of cortical astrocytes in awake mice expressing the calcium-sensitive lifetime sensor Tq-Ca-FLITS [25] in combination with implanted electrodes that allowed us to induce seizure-like events and record the electroencephalographic signal. Baseline activity was recorded for 3 minutes before inducing seizure-like events through brief electrical stimulation (Figure 4a), driving large calcium transients across the astrocytic network.

**Fig. 4.**
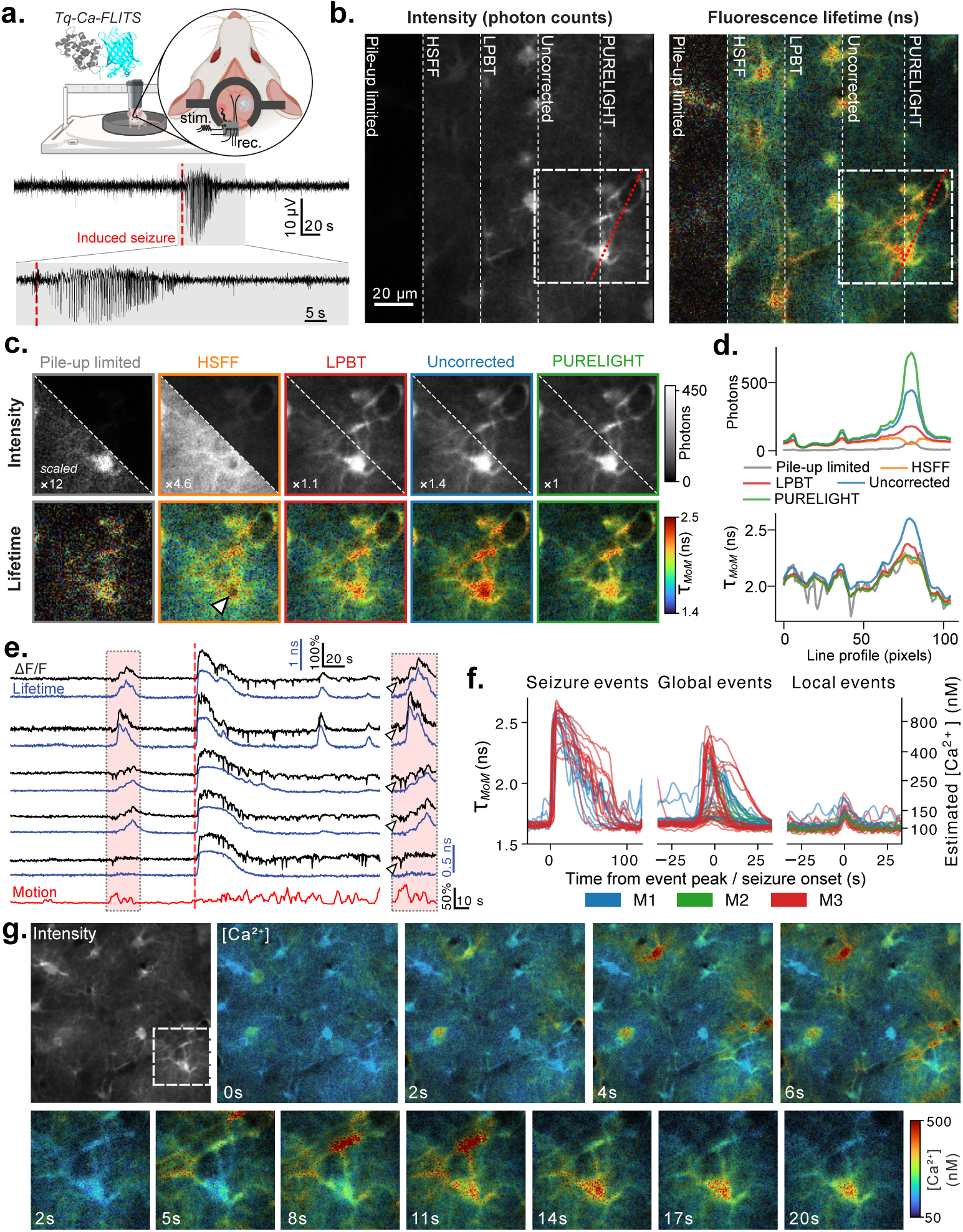
High-speed FLIM imaging of astrocytic calcium dynamics. **a)** Awake 2PM-FLIM imaging of astrocytes in the mouse somatosensory cortex (*∼*150 µm depth) expressing the Tq-Ca-FLITS sensor with simultaneous electroencephalogram and electrical stimulation induced seizure. **b)** Intensity and lifetime (*τ_MoM_*) images obtained from a single frame during the seizure event (0.34 s measurement time, 256 *×* 256 px; same FOV as panel g) processed by (from left to right): discarding photon counts to simulate a pile-up limited measurement (pile-up limited), keeping only excitation cycles with one photon detection (HSFF), applying an artificial blind time equal to the excitation period (LPBT), using all detected events (Uncorrected), applying the PURELIGHT correction (PURELIGHT). Dashed red line indicates where the profile analysis in (d) was performed. **c)** Enlarged view from panel b comparing the intensity and lifetime images obtained from each processing approach. Numbers in the bottom-left corner indicate the multiplier of photon counts applied for visualization for the bottom-left half of the image. **d)** Intensity and lifetime values along the line profile shown in red on panel b. **e)** Comparison of intensity Δ*F/F* (black) and mean lifetime (*τ_MoM_*) (blue) traces from multiple ROIs throughout the experiment. The red trace indicates brain motion as quantified by motion correction of intensity images. The shaded red box highlights a period of movement-induced astrocytic activity; artifacts induced by brain motion are visible as downward deflections in the intensity traces. The dotted line marks the time of seizure induction. **f)** Comparison in lifetime and calcium concentration of seizure (N=2 mice, n=21 ROIs), global (N=3, n=52) and local (N=3, n=26) events. **g)** Intensity-merged calcium concentration images showcasing spatiotemporal dynamics. *Top row:* Onset of a global calcium wave visualized at 2 s/frame. *Bottom row:* Close-up of the image above (white square) showing the propagation of calcium signals from endfeet wrapping around a blood vessel towards the soma, with subsequent persistent somatic activation.

For direct comparison, we processed a single image frame acquired during a seizure event with five different approaches: in addition to uncorrected data, HSFF, LPBT and PURELIGHT, we simulated a pile-up limited measurement by discarding an appropriate fraction of collected photons (Figure 4b-d, Supplementary Video 1). For both intensity and lifetime, the pile-up limited image displays a clearly insufficient SNR, showing the critical importance of pile-up correction. In the uncorrected data, bright somatic regions appear artificially longer-lived than dim processes, creating an intensity-lifetime correlation that could be mistaken for a genuine calcium gradient. Even though the HSFF correction provides undistorted lifetime values, the severe photon losses result in a noisy image with poor contrast (and even an “inversion” of the intensity profile in Figure 4d, as at very high count rates most excitation cycles contain more than one photon, and are thus discarded). LPBT and PURELIGHT are the only approaches capable of generating accurate intensity and lifetime images, but the latter clearly outperforms the former in terms of signal-to-noise ratio, resolving spatial variations in calcium concentration across soma, processes, and endfeet within a single sub-second frame.

To illustrate the advantages of lifetime-based calcium readouts we compared the temporal traces of intensity and lifetime from multiple astrocytic regions of interest (ROIs) (Figure 4e). We found that intensity traces were severely impacted by artifacts due to brain motion, while lifetime traces were largely unaffected (Figure 4e, Supplementary Figure 6a,b).

Calcium signals can arise both locally in individual astrocytes and globally across the entire astrocytic population [26]. To compare the calcium concentrations evoked under these two conditions, we combined our calibration data with the corrected life-time results to estimate the changes in calcium concentration for each event type (Methods), allowing us to directly compare responses across ROIs from experiments with several mice (Figure 4f). Global events reached peak concentrations of up to *∼*700 nM (mean 275 *±* 143 nM, *n* = 52 ROIs, N = 3 mice), whereas local events evoked transients under *∼*200 nM (mean 153 *±* 20 nM, *n* = 26, N = 3). Due to their robustness to motion, lifetime traces show a larger similarity between ROIs involved in the same global event compared to their intensity counterpart (Supplementary Figure 6c): a clear advantage for the identification and quantification of global events. The quantitative information on local and global events is also valuable to guide experiments with commonly used intensiometric sensors. In simulations of the response of GCaMP8m [27] and GCaMP6f [28] to the concentration changes observed in our experiments (Supplementary Figure 7), the former shows a higher occupancy at baseline and a larger relative amplitude of local events compared to global events, while the latter shows a larger Δ*F/F*_0_ for global events.

On the subcellular level, calcium signals exhibit a diversity of spatiotemporal dynamics [26, 29]. These events are challenging to quantify with conventional intensity-based imaging due to motion artifacts, scattering, and variable expression levels across subcellular compartments. Across our experiments, we observed a diversity of temporally resolved spatiotemporal events in individual astrocytes (Figure 4g, Supplementary Video 2). As an illustrative example, we report the observation of a transient initiated along blood vessels and propagating from the endfeet towards the soma (Figure 4g, bottom row). These observations demonstrate that PURELIGHT can be used to observe individual spatio-temporal events and to estimate the associated calcium concentrations on a subcellular level. In contrast, conventional TCSPC-based FLIM would struggle to capture these dynamics as the excitation power would need to be reduced to accommodate the brightest somatic regions, resulting in poor SNR in processes and endfeet.

To further test the limits of PURELIGHT in achieving ultrafast undistorted calcium concentration imaging in living animals, we imaged cortical neurons expressing Tq-Ca-FLITS at 30 Hz using resonant scanning (Figure 5a, Supplementary Video 3). As the imaging speed was about 10 times faster than in the astrocytic experiments, no method was able to provide a level of SNR sufficient to distinguish fine morphological features. To substantially improve the image quality, we applied unsupervised deep video denoising (UDVD, Figure 5b), revealing fine neuronal structures and lifetime contrast that would otherwise be buried in shot noise (Figure 5c). After denoising the HSFF and uncorrected images showed the same distortions observed in the astrocytic dynamics, with HSFF displaying a dramatic loss of spatial features and the uncorrected data displaying large lifetime errors in bright areas. PURELIGHT was the only method capable of generating accurate FLIM images at video-rate. To validate the combination of PURELIGHT and UDVD for retrieving accurate activity information, we compared ROI calcium traces obtained at different time-binning levels (Figure 5d). The PURELIGHT-UDVD trace at 30 Hz largely overlaps with the corresponding averaged trace (5 Hz), while significantly reducing variability.

**Fig. 5.**
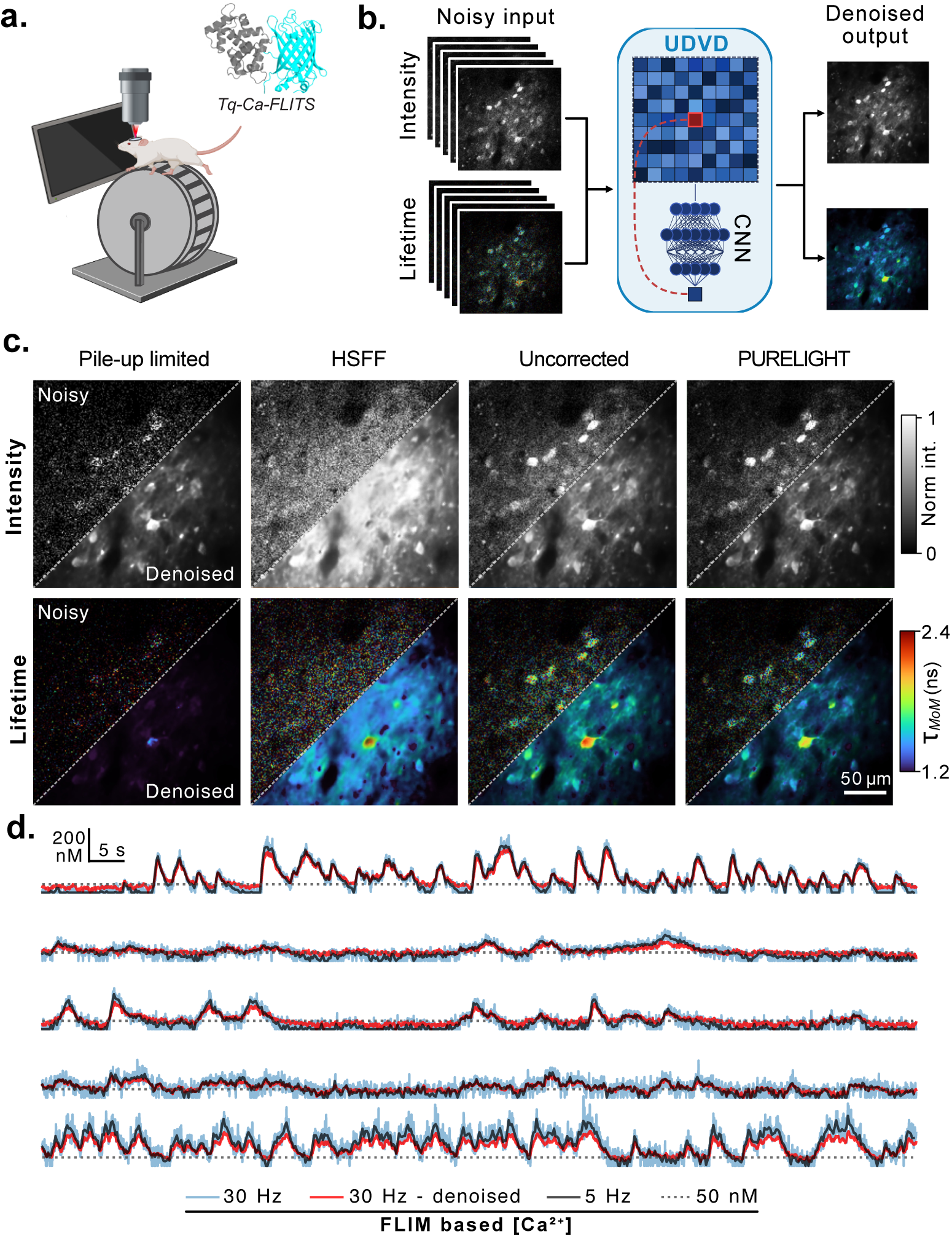
Combining PURELIGHT with self-supervised CNN-based denoising enables video-rate quantitative FLIM imaging of calcium activity in neurons. **a)** Awake 2PM-FLIM imaging of layer 2/3 neurons in the mouse visual cortex (*∼* 150 µm depth) expressing the Tq-Ca-FLITS sensor with simultaneous visual stimulation. **b)** UDVD-based [30] self-supervised denoising pipeline for FLIM images. The noisy intensity and lifetime (e.g. *τ_MoM_*) image series are combined to train a two-channel 3D convolutional neural network (CNN) to predict held-out pixel values based on their spatiotemporal neighborhood. After training, the network weights are fixed and the whole two-channel image sequence is predicted to obtain denoised intensity and average lifetime image series. **c)** Comparison of noisy (top left) and denoised (bottom right) images for different pile-up correction methods. **d)** Calcium concentration traces of manually selected cell body ROIs (n=5 ROIs, N=1 mouse). The light blue line shows the concentration as computed from the mean lifetime of the ROI for each noisy frame acquired at 30 Hz, the red line is computed from the denoised 30 Hz images, and the black line is computed from a 6-frame average of the lifetime images, reducing sampling rate to 5 Hz. The dotted line shows the 50 nM level as an absolute reference for each ROI.

### 2.5 Undistorted TCSPC improves SNR and eliminates analog detection artifacts in temporally multiplexed dual-area calcium imaging

The study of inter-area communication, functional connectivity, or the coordinated dynamics of distributed circuits requires the simultaneous observation of neuronal activity across spatially distant brain regions. Dual-area two-photon imaging through temporal multiplexing provides an elegant solution by interleaving excitation pulses in time so that multiple fields of view (FOVs) can be imaged simultaneously without compromising spatial or temporal resolution [31, 32] (Figure 6a). With conventional analog integration, the pixel value for each area is computed by integrating the detector signal within the corresponding time window. Critically, the analog baseline drifts with the average photon rate, so that a bright event in one area shifts the baseline of the opposite area (Figure 6b), creating an inverted artifact that can dominate the signal in dim regions. In contrast, photon counting detects threshold crossing and assigns the photons to the correct area based on their measured arrival time, effectively eliminating baseline crosstalk. When no lifetime information is needed, our XTTTR format allows us to implement the simpler multi-photon counting (MPC) approach (Figure 6b), which can be used to obtain accurate intensity even in the presence of very significant pile-up (Supplementary Figures 3,4).

**Fig. 6.**
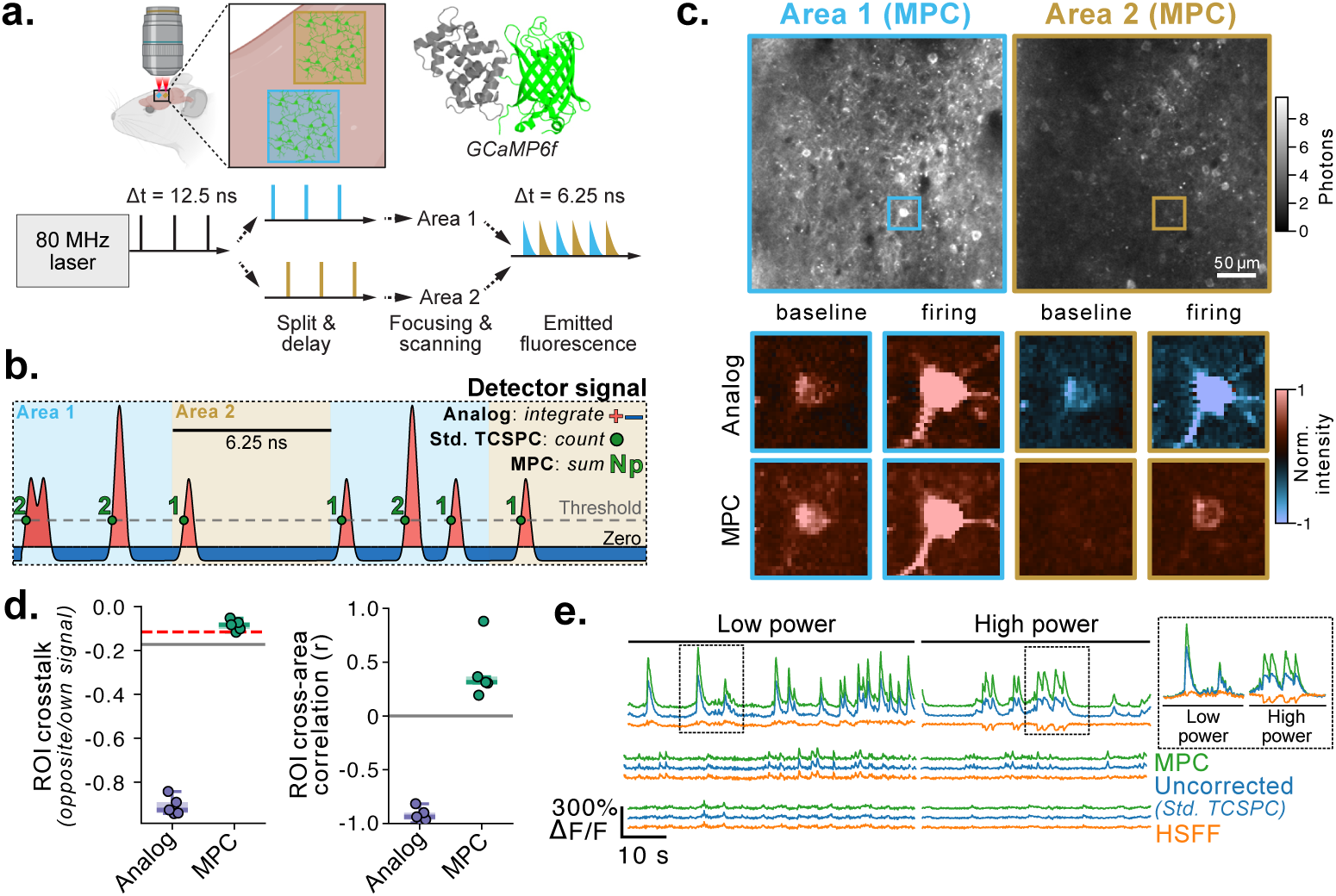
TCSPC enables temporally multiplexed dual-area calcium imaging with accurate intensity readouts. **a)** Working principle of temporally multiplexed dual-area imaging. The output of an 80 MHz pulsed laser is split by a 50/50 beam splitter and one path is delayed by half the interpulse interval (6.25 ns) such that each FOV is excited by alternating pulses at the full 80 MHz rate. The fluorescence from both areas reaches the same detector, and the two signals must be separated based on their arrival time relative to the excitation pulses. **b)** Determination of pixel intensity from the detector signal using either analog integration, standard TCSPC or MPC-based detection: analog integration sums the full signal including the drifting baseline (negative areas, blue), whereas standard TCSPC counts threshold crossings (green circles) within defined time gates. Multi-photon counting (MPC) builds on top of standard TCSPC by assigning a number of photons (*N_p_*) to each threshold crossing, estimating the true number of photons arriving at the detector (Supplementary Figures 3,4d). **c)** Simultaneous two-photon calcium imaging (GCaMP6f) of two cortical areas (visual cortex, layer 2/3) during baseline and neuronal firing at low and high excitation power. Close-up images show a region with a bright cell body in area 1 and no cell bodies in area 2. Top: analog integration intensity images. Bottom: MPC intensity images. **d)** Crosstalk quantification. Left: ratio of the signal measured in the opposite area to the signal in the own area for each ROI. Right: Pearson correlation between the same ROI measured in the two areas. Dashed red line: expected residual crosstalk from a 2.6 ns fluorescence lifetime. The analog images showed a median crosstalk of *−*90.8% (median r = *−*0.94) while the TCSPC images exhibited only 10.7% median residual crosstalk (median r = 0.31), consistent with the expected leakage of *∼*7% from the fluorescence tail of GCaMP (*τ ≈* 2.6 ns). **e)** Δ*F/F*_0_ traces for three manually segmented ROIs from two measurements of the same FOV with different excitation powers. The XTTTR data was processed with different pile-up correction methods (HSFF: orange, Uncorrected - Std. TCSPC: blue, MPC: green).

We performed simultaneous two-photon calcium imaging of two cortical areas in awake mice expressing GCaMP6f in layer 2/3 neurons and compared the MPC results with conventional analog integration (Figure 6c,d, Supplementary Video 4). The superiority of MPC can be observed in the close-up images in Figure 6c: when a bright cell fires in the first area, the corresponding region in the second area shows only a small positive crosstalk, whereas with analog detection a strong inverted signal is observed. Beyond removing crosstalk, combining TCSPC with the PURELIGHT or MPC corrections also recovers the intensity linearity that is lost with traditional photon counting at high count rates. The HSFF correction severely limits the detected count rate, reaching the point of negative Δ*F/F*_0_ values at higher excitation powers, while uncorrected photon counts result in a diminished dynamic range where higher excitation powers lead to lower Δ*F/F*_0_ values (Figure 6e).

## 3 Discussion

In this work, we introduced PURELIGHT, a TCSPC acquisition and correction framework that preserves both photon counts and fluorescence decays under conditions that produce substantial pile-up. By combining fast digitization with an extended time-tagged data format, PURELIGHT records not only photon arrival times but also pulse-shape information needed to estimate detector active time and timing shifts. Allowing intensity and lifetime distortions to be corrected from the same photon stream, PURELIGHT enables video-rate FLIM and quantitative intensity imaging across commonly used detector classes.

Most previous strategies for avoiding pile-up fall short of the requirements for seamlessly transitioning two-photon microscopes from analog integration to TCSPC detection. Photon rejection [11, 33], imposed long dead times [12, 34] and conservative count-rate operation can recover undistorted temporal distributions, but at the cost of photon efficiency and/or intensity linearity. Model-based correction can account for decay distortions, but constrains downstream analysis to computationally intensive and noise-sensitive exponential fitting[13]. Active-time correction provides a more direct route to recovering intensity and lifetime, but remains sensitive to detector pulse shape and time walk. Fast digitization of photodetector signals offers an alternative route to time-domain FLIM and has been used to directly sample amplified detector waveforms [35–38]. However, direct waveform sampling typically trades TCSPC timing precision and data efficiency for high-throughput acquisition, resulting in reduced temporal resolution and increased noise at low photon rates [39]. A conceptually similar method, computational photon counting, recovers discrete photon events from digitized waveforms and discriminates multi-photon signals [40]. However, no pile-up correction was applied and the transfer and processing of raw ADC data on the PC poses a significant bottleneck.

PURELIGHT can be considered a hybrid between active-time correction and fast digitization methods. By digitizing the detector response and extracting additional pulse parameters directly on the FPGA, PURELIGHT extends the active-time correction method with time-walk compensation, making the method effectively detector-agnostic. Moreover, XTTTR data acquisition maintains the precision and data efficiency of TCSPC-based detection at low count rates while enabling accurate measurements and a variety of pile-up correction approaches at high count rates.

This versatility is important for two-photon microscopy because the limitations of analog integration and conventional TCSPC occur in complementary regimes. At low photon budgets, analog PMT integration introduces excess variance from pulse-height fluctuations and electronics, whereas photon counting preserves the statistics of photon emission. At high photon rates, conventional TCSPC loses quantitative accuracy because of pile-up. PURELIGHT removes this constraint, allowing us to fully exploit the benefits of photon-resolved detection.

We demonstrated the advantages of undistorted TCSPC detection across a wide range of biologically relevant scenarios and signal strengths. In the high count rates regime, we performed accurate two-photon FLIM imaging at over 160 Mcps peak count rate with both HPDs and SiPMs, resolving the spatiotemporal dynamics of calcium concentrations in astrocytes and neurons with subcellular resolution at video rate. In temporally multiplexed measurements, we showed how time-gated multi-photon counting removes analog crosstalk between the channels while recovering the dynamic range and intensity linearity lost with traditional photon counting. In photon starved conditions, we have proven how TCSPC detection significantly improves signal-to-noise ratio in ratiometric autofluorescence imaging with PMTs, with immediate applicability to label-free classification and diagnostics using standard detectors available in the vast majority of microscopes. Finally, as a proof of the unique capabilities enabled by the simultaneous correction of intensity and lifetime, we have demonstrated a novel approach in which ratiometric readouts are internally calibrated using FLIM, combining high signal-to-noise ratio with quantitative precision.

The practical limitations of PURELIGHT are relatively minor and technical rather than fundamental. First, the current post-processing implementation prevents accurate real-time visualization and increases the computational cost for large datasets. Second, XTTTR acquisition requires higher data throughput than conventional TTTR. Although substantially more efficient than continuous waveform streaming, XTTTR transmission can become limiting when very high count rates are collected for extended time periods. Third, the usable count rate range remains detector-dependent. Detectors with broad single-photon responses tag fewer threshold crossings and require longer additional blind times to avoid ambiguous time tagging, reducing signal-to-noise ratio. Faster detectors will directly extend the usable count-rate range, whereas integration of photon detection with dedicated computational hardware should reduce data-transfer and post-processing bottlenecks and enable real-time correction.

Most importantly, PURELIGHT unifies intensity imaging and FLIM into a single acquisition, allowing a laser-scanning microscope to operate entirely in photon-counting mode with no loss of speed, dynamic range or accuracy. A single acquisition with PURELIGHT yields information that previously required separate experiments or distinct hardware configurations, such as shot-noise-limited intensity, undistorted fluorescence lifetime, and timing information for temporal multiplexing. Because the approach is implemented as a detector-agnostic digitization and processing pipeline, it can be easily integrated into existing microscopes by employing detection electronics capable of providing an XTTTR data stream to be processed with our open-source pile-up correction algorithms. We anticipate that making the complete photon-level information routinely available at video rate will expand the practical use of FLIM, FRET, temporal multiplexing, and other time-resolved techniques from specialized applications to standard tools across the broader imaging community.

## 4 Methods

### 4.1 Custom ADC-based TCSPC electronics

A custom TCSPC system was built around a KRM-4ZU47DR module (Knowledge Resources GmbH, Switzerland), based on a Gen 3 AMD RFSoC XCZU47DR. The module provides eight high-speed ADC channels operated simultaneously at 5 GS*/*s. One channel was reserved for the laser synchronization signal, while the remaining seven channels were available for detector signals from PMTs, HPDs, SiPMs, or similar photosensors.

All enabled detector channels were continuously monitored for crossings of a user-defined threshold, mimicking a time-to-digital converter architecture [12]. When a threshold crossing was detected, an event was generated and ADC samples around the crossing were retained for on-FPGA processing. These samples were used to extract the XTTTR event information (Figure 1). The rising or falling edge was linearly interpolated into 16 segments, improving the timestamp precision from the ADC sampling interval of 200 ps to 12.5 ps.

For the laser synchronization channel, the FPGA can lock to a periodic signal up to 125 MHz. Rather than assigning an independent timestamp to each laser period, 1,024 threshold crossings were recorded and used to calculate the timing of individual synchronization events. This provided a more precise time reference than single-trigger timestamping. The same block also reported the detected trigger frequency and flagged synchronization errors when the measured and calculated trigger times differed by more than a user-defined tolerance.

In addition to the analog detector and synchronization channels, the system provides 16 digital inputs timestamped at the internal clock resolution of 4 ns. Two of these inputs were used for the microscope frame and line clock signals required for image reconstruction. Analog and digital events were sorted by timestamp at a maximum rate of one event per internal clock cycle, corresponding to 250 MHz. The relative photon arrival times (*t*) were then computed and the events were assembled into the selected output format. Depending on the acquisition settings, the data contains the detection channel, absolute time (*T*), and relative time (*t*), with an optional additional 16–32 bits per event encoding pulse-shape parameters in the XTTTR format. The resulting data stream can reach a maximum rate of 16 Gbit*/*s.

The event stream was buffered in local DDR memory and forwarded to a separate FPGA block for real-time image generation. This block computed intensity images from the number of events per pixel and “fast FLIM” images from the average relative arrival time, enabling live visualization. These preview images were also buffered in local DDR memory.

Both DDR-buffered streams were read by the local processor system and transmitted to the host computer through a 10 Gbit*/*s TCP/IP connection. A block diagram of the FPGA implementation is shown in Supplementary Figure 1.

### 4.2 Custom SiPM Module

A custom SiPM detector was built around a Hamamatsu S13362-3050DG MPPC [41]. The detector consists of a square SiPM with an active area of 9 mm^2^, mounted in a sealed barrel package with a thermoelectric cooler (TEC) on the back side. The module provides the SiPM bias voltage and regulates the TEC to maintain the detector at *−*10 *^◦^*C.

The SiPM output current was converted to voltage using a 120 Ω resistor and amplified by two non-inverting amplifier stages based on the TI OPA855 [42]. The two stages provided a combined gain of 300 and used pole-zero feedback compensation to flatten the frequency response from DC to approximately 400 MHz. The single-photon response of the SiPM module is shown in Figure 2a.

### 4.3 TCSPC simulation

To evaluate the performance of the correction methods, we extended a previously implemented Monte Carlo TCSPC simulation [12] to reproduce the XTTTR acquisition with the ADC-based TCSPC electronics described above. The simulated threshold-crossing detection was modified to record the additional pulse parameters required for XTTTR processing. The correction methods were then applied to the simulated XTTTR data using the same processing pipeline as for the experimental measurements.

### 4.4 Collection and processing of XTTTR data

For all experiments, TCSPC data were acquired in the 64-bit XTTTR format using a C++ acquisition library controlled from a Python-based graphical interface. During acquisition, a real-time preview stream containing intensity and fast-FLIM images was displayed directly in the GUI for live visualization. In parallel, the full XTTTR event stream was handled by the C++ library, which controlled acquisition completion and wrote the data to disk.

The stored XTTTR data were processed offline to generate corrected intensity and FLIM images. First, the relevant correction procedure was applied to the event stream. The corrected events were then assigned to image pixels using the recorded frame and line markers. For PURELIGHT correction, each event contributed its pre-computed weight to the photon-count histogram instead of a unitary count. For the HSFF and LPBT corrections, only events retained after the corresponding filtering step were included. For multi-photon-counting intensity images, the estimated number of photons associated with all events within each pixel was summed and reported as the total photon count.

### 4.5 Multi-photon counting (MPC)

Using the pulse width (*t_p_*) and height (*H_p_*) available in the XTTTR format, we can visualize a 2D heatmap of the distribution of all detected events at high count rates with various sample lifetimes (Supplementary Figures 3-5d). Both the HPD and SiPM heatmaps present prominent clusters of detection events with different widths and heights. Using these high count rate short and long lifetime measurements parallel linear boundaries can be drawn between these clusters, starting from the cluster of single photon events (*t_p_* and *H_p_* in the SPR range of the detector) and assigning each successive cluster one additional photon. An additional boundary can be included to reject detection events deemed too small to be photon detections (0 photons). After calibration, MPC assigns a number of photons to each detection event based on its pulse width and height and the predefined boundaries. Then, intensity images are assembled by summing the number of photons in all of the events for each pixel. Supplementary Figures 3-5c show the linearity of the MPC count rate across different sample lifetimes with all detectors.

### 4.6 2P microscopy setup

All *in vivo* and *ex vivo* imaging was performed on custom-built two-photon microscopes equipped with our ADC-based TCSPC electronics.

For astrocyte calcium, ATP FRET, and optic nerve autofluorescence imaging, a tunable-wavelength 80 MHz femtosecond laser (Chameleon Discovery NX with Total Power Control, Coherent, Saxonburg, PA, USA) was sent into a custom two-photon microscope [43] and focused onto the sample through a water immersion objective (16*×* NA 0.8, Nikon)/(W Plan-Apochromat 20*×* NA 1.0, Zeiss, Jena, Germany)/(25*×* NA 1.05 XLPLN25XWMP2, Olympus). The fluorescence was collected through the same objective and split from the excitation light using a dichroic mirror (KS93 Cold Light Mirror, 685 nm LP, QiOptiq, Rhyl, UK). The collected light was then divided by wavelength using a set of dichroic mirrors (F38-560, F38-506; AHF Analysentechnik, Tübingen, Germany) and bandpass filters (Brightline HC 542/50 or Brightline HC 475/50, each combined with 785SP Edge; Semrock, Rochester, NY, USA). For the astrocyte calcium experiments the fluorescence was detected using a hybrid photomultiplier detector (PMA Hybrid 40 mod, PicoQuant, Berlin, Germany) in combination with a tightly focusing lens (LB1761-A, Thorlabs, Newton, NJ, USA) to minimize back-reflections. For the ATP FRET experiments two of our custom 3*×*3 mm^2^ SiPM were used to collect both emission channels simultaneously. For the optic nerve autofluorescence experiments two GaAsP PMTs (H10770PA-40, Hamamatsu, Japan) were used. The output of each PMT was amplified by a TA1800B pulse preamplifier (FAST ComTec, Germany) and their gain was set such that the average peak voltage from the SPR was around 70 mV after amplification.

For neuronal GCaMP and Tq-Ca-FLITS imaging, a separate dual-area two-photon microscope [44] was used, equipped with a Titanium-sapphire laser (MaiTai HP DeepSee, Spectra-Physics) at 80 MHz and a 16*×* NA 0.8 water-dipping objective (Nikon). Imaging was performed on one or both areas at the same time, with the fluorescence collected through the same objective and split from the excitation light using a dichroic mirror. The collected light was then filtered to remove residual excitation light (542/50 nm, Semrock) and detected using a Hamamatsu hybrid photodetector (R11322U-40, Hamamatsu, Japan). The spatiotemporal multiplexing scheme was demultiplexed either by using virtual analog channels in ScanImage Premium 2023 (MBF Bioscience, Williston, VT, USA) or using the XTTTR data from the TCSPC electronics.

### 4.7 Imaging of fluorescent solutions and instrument response function determination

To evaluate the system under controlled conditions, we used sodium fluorescein solutions at varying concentrations in pH 8 phosphate-buffered saline. Unquenched fluorescein was prepared at a concentration of 5 *×* 10*^−^*^6^ M. To obtain shorter monoexponential lifetimes, fluorescein was quenched with potassium iodide (KI) at several concentrations up to 0.72 M, yielding lifetimes ranging from *∼*0.6 ns to 4 ns. For quenched solutions, the fluorescein concentration was increased to 5 *×* 10*^−^*^5^ M to compensate for the reduction in fluorescence quantum yield.

Imaging was performed at 950 nm excitation using a 20*×* water immersion objective (W Plan-Apochromat 20*×*/1.0, Zeiss, Jena, Germany). To obtain measurements at different count rates, the excitation power was varied using the laser’s integrated acousto-optic modulator. A beam sampler (BSF10-B, Thorlabs) placed in the optical path directed a small fraction of the beam to a photodiode (PDA50B2, Thorlabs), which was calibrated against a power meter (S175C, Thorlabs) placed under the objective to allow continuous monitoring of the power at the sample during FLIM measurements. For each excitation power, data were collected in the XTTTR format and processed using all correction methods. Each measurement was repeated with the HPD, SiPM, and PMT detectors. A single decay per image was computed by subtracting a background image and summing over all pixel-associated decays, ensuring *>* 10^4^ photons per decay.

Instrument response function (IRF) traces were measured using second harmonic generation (SHG) from KH_2_PO_4_ crystals at 1040 nm. Background images were acquired using a pure water solution and subtracted from the fluorescein and SHG data when performing lifetime analysis.

### 4.8 Fluorescence lifetime estimation methods

Three lifetime estimators are used throughout this work, and are denoted *τ_fit_*, *τ_I_* and *τ_MoM_*. *τ_fit_*is the lifetime of a monoexponential fit (*n* = 1), used for the fluorescent solutions and the IRF characterization, where the sample is known to be monoexponential. *τ_I_*is the intensity-weighted mean lifetime of a biexponential fit (*n* = 2), used for the genetically encoded FRET and lifetime sensors, whose decays are not monoexponential because they contain donor populations in different conformational or binding states:

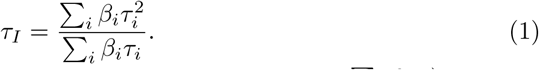

The intensity weighting (rather than the amplitude-weighted mean ∑*_i_ β_i_τ_i_*) was chosen because *τ_I_* is proportional to the steady-state fluorescence intensity of the sensor and is therefore the quantity that is directly comparable to intensity-based ratiometric readouts. Fitted lifetimes (*τ_fit_* and *τ_I_*) were obtained by iterative reconvolution using the FLIMfit library [45], accessed through the Python interface of our acquisition and analysis library. In both cases the lifetimes and their amplitudes were free parameters, initialized at 3 ns and 1 ns, and the fit was performed globally across an image. Any pixels with non-physical parameters (e.g. negative lifetimes or amplitudes) were excluded from downstream analysis.

*τ_MoM_* is a fit-free estimate of the mean arrival time obtained by the method of moments [46], i.e. the center of mass of the decay histogram. Since the first moment is measured with respect to an arbitrary time origin, the decays were first shifted so that the start of the rising edge of the decay coincides with *t* = 0. While this does not account for the incomplete decay of longer lifetimes it minimizes the wrap-around contribution to the moment. The lifetime is then

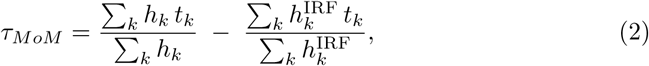

where the second term is the first moment of the IRF, subjected to the same alignment procedure, which removes the contribution of the finite width and position of the excitation and detection response.

*τ_MoM_* requires no fitting, no initial guesses and no per-pixel convergence, and it is statistically efficient: for an ideal monoexponential decay fully contained within the period, the mean arrival time is the maximum-likelihood estimator of the lifetime. It is therefore the estimator used for the per-pixel, per-frame lifetime images of the video-rate *in vivo* recordings, where the number of photons per pixel and frame is too low for reliable fitting, as well as for the calcium concentration estimates described below.

### 4.9 Estimation of calcium concentrations

Intracellular calcium concentrations were estimated from the corrected fluorescence lifetime images on a per-pixel basis. The CaFLITS sensor reports calcium binding through an increase in its fluorescence lifetime that follows a Hill relationship, so the mean lifetime *τ_MoM_* maps onto the fractional sensor occupancy *f* between a calcium-free lifetime *τ*_min_ and a calcium-saturated lifetime *τ*_max_:

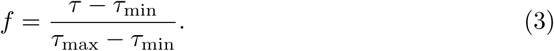

Inverting the Hill equation then yields the calcium concentration

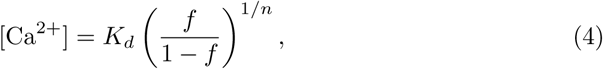

using the CaFLITS in-cell affinity *K_d_* = 265 nM and Hill coefficient *n* = 1.63[25]. Before inversion, *f* was clamped to the range [10*^−^*^4^, 1*−*10*^−^*^4^] to keep the concentration finite at the extremes of the sensor’s dynamic range.

The lifetime bounds were calibrated from the experimental data. The saturated lifetime *τ*_max_ = 2.69 ns was taken as the mean of the peak lifetimes reached during a set of near-saturating seizure events, where astrocytic calcium is expected to plateau at the top of the sensor’s range. The calcium-free bound *τ*_min_ = 1.41 ns was set such that the median lifetime across baseline recordings (1.63 ns) corresponds to 100 nM, a calcium concentration that is consistent with reported resting astrocytic levels [47]. The same calibration (*τ*_min_, *τ*_max_, *K_d_*, *n*) was applied to all recordings, enabling direct comparison of calcium concentrations across ROIs, cell types, and animals.

### 4.10 Unsupervised time-series denoising

For time-series denoising we processed the intensity and lifetime images data using a custom implementation of the Unsupervised Deep Video Denoising network (UDVD, [30]). The blind-spot network is the original architecture, configured here with a 9-frame temporal receptive field. Each acquisition channel (intensity and lifetime) was normalized independently to unit mean and denoised jointly by a single model making use of the multi-channel setup originally intended for RGB (3-channel) images. Unlike the reference UDVD implementation, which models a per-pixel Gaussian prior (mean + covariance) and forms the denoised estimate as the Bayesian posterior mean fusing the prediction with the noisy observation under a single-*σ* Gaussian noise model, we minimized a plain mean-squared blind-spot loss. Because our data are photon-limited (Poisson, signal-dependent variance) and include a fluorescence lifetime channel, the single-*σ* Gaussian noise assumption underlying the likelihood loss is not satisfied; the MSE objective makes no noise-model assumption and yields the conditional mean of the clean signal. This choice trades a modest loss of high-frequency detail for robustness and avoids discretization artifacts at low photon counts. Training was fully self-supervised (no clean targets, no synthetic noise) over 128*×*128 patches (stride 64), 80 epochs, Adam (learning rate 1 *×* 10*^−^*^4^). 4-way data augmentation (horizontal, vertical, and combined flips + original) was used in cases where the number of training patches was *<* 20, 000.

### 4.11 Animals

All experimental procedures were approved by the local veterinary authorities in Zurich and were conducted in accordance with the guidelines of the Swiss Animal Protection Law (Veterinary Office, Canton of Zurich; Act of Animal Protection, 16 December 2005, and Animal Protection Ordinance, 23 April 2008).

### 4.12 Surgical interventions

To investigate astrocytic calcium dynamics during seizures, simultaneous electrical and optical recordings were performed in awake mice. Eight- to fourteen-month-old C57BL/6 mice were used for the experiments.

#### Surgical procedures

Animals were deeply anesthetized by intraperitoneal injection of a mixture of fentanyl (0.05 mg/kg body weight (BW); Sintenyl, Sintetica), midazolam (5 mg/kg BW; Roche), and medetomidine (0.5 mg/kg BW; Orion Pharma). Anesthesia was supplemented as needed, approximately 60 min after induction. Once deeply anesthetized, animals were placed in a stereotaxic frame (Model 900; David Kopf Instruments). Vitamin A eye ointment was applied to prevent corneal desiccation. To prevent dehydration, prewarmed Ringerfundin was injected subcutaneously before the start of the procedure at a dose of 10 mL/kg BW. Oxygen was delivered at 200 mL/min to prevent hypoxemia. Vital signs and body temperature were continuously monitored and maintained using a MARTA Pad system (Vigilitech).

#### Head plate implantation

The scalp was shaved, disinfected with Kodan (Schülke & Mayr), and locally anesthetized by subcutaneous injection of a mixture of lidocaine (10 mg/mL) and bupivacaine (5 mg/mL). The skull was exposed through a midline incision approximately 1.2–1.5 cm in length. Connective tissue was carefully removed, and the skull surface was cleaned and coated with a bonding agent (One Coat 7 Universal, Coltene). A custom-made stainless steel head plate was positioned centrally over the exposed skull and fixed in place using multiple layers of blue light-curing dental cement (Tetric EvoFlow, Ivoclar Vivadent), while leaving the regions designated for electrode implantation and craniotomy exposed.

#### Electrode fabrication and implantation

Custom electrodes were fabricated from PFA-coated stainless steel wire with a diameter of 0.008 inches. Bipolar stimulating electrodes were prepared by twisting two insulated wires together and soldering them to a common connector (stiftleiste female 50). The stimulating electrode was implanted into the left ventral CA1 region of the hippocampus (3 mm anteroposterior, 3 mm mediolateral, and 3 mm dorsoventral relative to bregma). Bilateral cortical recording electrodes and a cerebellar reference electrode were subsequently implanted. All electrodes were secured to the skull using dental cement.

#### Craniotomy and AAV injection

A craniotomy was performed over the right somatosensory cortex using a dental drill with a 0.2 mm diameter burr (H-4-002HP, Rotatec GmbH). Following craniotomy, adeno-associated virus injections were performed using a custom-made microinjector. AAV9-GFAP-CaFLITS (1.7 *×* 10^13^ vg/mL; Viral Vector Facility, University of Zurich, Zurich, Switzerland) was injected through a glass capillary at two depths below the brain surface, 350 *µ*m and 200 *µ*m, with 100 nL delivered per injection. Large blood vessels were avoided to minimize the risk of bleeding. After viral injection, a transparent glass coverslip was gently placed over the exposed brain region and sealed with light-curing dental cement (Tetric EvoFlow, Ivoclar Vivadent) to create a chronic cranial window. At the end of the procedure, anesthesia was reversed by intraperitoneal injection of a mixture of flumazenil (0.9 mg/kg BW) and atipamezole (45 mg/kg BW). Postoperative analgesia consisted of buprenorphine (0.1 mg/kg BW, subcutaneous) and carprofen (10 mg/kg BW, subcutaneous). Carprofen treatment was continued twice daily for three days after surgery.

### 4.13 *In vivo* imaging

#### Behavior training for awake two-photon imaging

After recovery, animals were habituated to head fixation by restraining them using the implanted head plate several times per day. Restraint duration was gradually increased from a few seconds to several minutes. Additional training sessions were performed in the microscope setup, with the microscope and shutters operating, to familiarize the animals with the experimental environment. After a training period of approximately two weeks, mice tolerated head fixation during imaging sessions while freely moving on a custom-made air-lifted platform or running wheel.

#### Electroencephalographic recordings and seizure induction protocol

Simultaneous electroencephalographic (EEG) recordings were acquired throughout each imaging session. Non-convulsive electrographic seizures were elicited by electrical stimulation of the ventral hippocampus through the chronically implanted bipolar electrode. Cortical activity was monitored using the implanted cortical recording electrodes. The after-discharge threshold was determined individually for each animal using 2 s stimulation trains with a minimum amplitude of 20 *µ*A at 50 Hz, delivered with a stimulus generator (STG3000 series, Multi Channel Systems). Seizures were identified by an experienced researcher and defined as spike–wave discharges exceeding two times the baseline amplitude, with a frequency greater than 5 Hz and a duration longer than 20 s. EEG signals were acquired using a Multi Channel Systems recording system, enabling precise synchronization between calcium imaging and electrophysiological datasets. Throughout each imaging session, mice were continuously observed and video-recorded with an infrared camera using custom-written PyAwake software.

### 4.14 *Ex vivo* acute brain slice imaging

For acute brain slice imaging, ten-week-old transgenic Thy1-ATP mice [48] were used. Mice were deeply anesthetized with isoflurane and decapitated, following which the brain was dissected in ice-cold cutting solution (in mM: 135 NMDG, 1 KCl, 1.2 KH_2_PO_4_, 1.5 MgCl_2_, 0.5 CaCl_2_, 10 glucose, 20 choline bicarbonate; pH 7.4). Coronal sections (300 µm) were cut with a vibratome (Vibration microtome, HM 650V, VWR) in the cutting solution. Slices were then transferred to a recovery bath containing ACSF (in mM: 120 NaCl, 2.5 KCl, 1.25 NaH_2_PO_4_, 25 NaHCO_3_, 1 MgCl_2_, 2 CaCl_2_, 10 glucose, 10 sucrose, 1 sodium lactate, 0.1 sodium pyruvate; pH 7.4), kept at 34 *^◦^*C for half an hour and then kept at room temperature until imaging. All solutions were continuously bubbled with 95 % O_2_ and 5 % CO_2_. For imaging, slices were transferred to a recording chamber (RC26, Warner Instruments, Hamden, Connecticut, United States) mounted on a temperature-controlled Mini Bath Chamber (Luigs & Neumann, Ratingen, Germany) through an aluminum adapter and continuously perfused with oxygenated ACSF at a flow rate of 3 mL*/*min using a pumping system based on the PiFlow system [49]. Throughout all experiments, the temperature was maintained at 34 *^◦^*C. Slices were imaged either in pure ACSF or with the addition of 5 mM sodium azide (NaN_3_) as necessary.

### 4.15 *Ex vivo* optic nerve imaging

Acute optic nerve preparations for two-photon imaging were prepared as previously described [50]. Following isoflurane anesthesia and decapitation, optic nerves were dissected and transferred to an interface perfusion chamber continuously supplied with ACSF containing 126 mM NaCl, 3 mM KCl, 2 mM CaCl_2_, 1.25 mM NaH_2_PO_4_, 26 mM NaHCO_3_, 2 mM MgSO_4_ and 10 mM glucose (pH 7.4). The ACSF was maintained at 37 *^◦^*C and continuously equilibrated with 95% O_2_ and 5% CO_2_. The ends of each nerve were inserted into custom-made suction electrodes filled with ACSF. The preparation was mounted on a custom-built two-photon microscope equipped with a Chameleon Ultra II Ti:sapphire laser (Coherent) and a 25× water-immersion objective (XLPLN 25×/1.05 WMP2, Olympus).

### 4.16 Astrocytic calcium event analysis

Event traces and concentration statistics (Figure 4f) were derived from PURELIGHT-corrected recordings of three mice, comprising three baseline and two seizure recordings acquired at 3 Hz (256 *×* 256 px, 1080 frames, 360 s each).

Rigid and piecewise-rigid motion correction was performed with NoRMCorre [51] through the CaImAn Python interface [52], in two successive passes of rigid followed by piecewise-rigid registration with maximum shifts of 18 and 5 px, a 32 *×* 32 px patch grid, 16 px overlap and a maximum patch deviation of 3 px. Since the fluorescence intensity of Tq-Ca-FLITS is largely calcium-independent, the transformations were estimated on the intensity channel and applied unchanged to the lifetime images.

Somatic ROIs were segmented manually from the motion-corrected intensity images (10–12 per recording, 54 in total). Local-event ROIs were drawn manually on lifetime maps smoothed with a Gaussian kernel of *σ* = 2 in time and space (*n* = 27; M1: 7, M2: 3, M3: 17). Lifetime traces were computed as the mean *τ_MoM_* over the pixels of each ROI, excluding non-positive values, after smoothing the lifetime movies along time with a Gaussian kernel of *σ* = 2 frames (0.67 s).

Three event classes were analysed. *Seizure events* comprise every soma of the two seizure recordings over a fixed window 150 s to 300 s after recording onset (*n* = 22). *Global events* were detected on the mean lifetime across all somata of a recording during the 180 s baseline period, as excursions above the median plus two standard deviations lasting at least six consecutive frames, and extracted for every soma over *±*100 frames (*±*33 s) around the midpoint of the supra-threshold interval (5 events, *n* = 54 soma-event traces). *Local events* are the manually defined ROIs above, extracted over the same window centred on the frame each was drawn on.

Lifetimes were converted to calcium concentrations per trace using *τ*_min_ = 1.41 ns and *τ*_max_ = 2.69 ns, always before averaging, as the Hill relation is non-linear.

## 5 Code and data availability

The C++ acquisition library, Python-based acquisition GUI, and Python-based correction algorithms are open source and will be available at https://gitlab.uzh.ch/einlab/purelight.

## Acknowledgements

This work was partially funded by the Innosuisse Innovation Agency (Grant No. 52207.1). P.R. was supported by a grant from the Swiss National Science Foundation (Ambizione grant PZ00P3 209114).

