## Supplementary Information for "PURELIGHT: a quantitative photon-counting framework unifying intensity and lifetime imaging at video rate across detector technologies"

Felipe Velasquez Moros *et al.*

**Supplementary Figures** **10**

### List of supplementary videos

1. **Astrocytic $Ca^{2+}$ method comparison**: FLIM video of the data shown in Figure 4b-d (baseline and seizure) comparing the different correction methods.

2. **Baseline astrocyte $Ca^{2+}$**: FLIM video of the data shown in Figure 4g and other local and global events during baseline with a narrower color scale (1.4 ns-2.2 ns) to improve visualization of smaller events.

3. **Video rate neuronal $Ca^{2+}$ FLIM**: Noisy and denoised 30 Hz recording of neurons with Tq-Ca-FLITS.

4. **Dual area calcium imaging analog vs TCSPC comparison**: Two GCaMP6f intensity channels of different cortical areas recorded simultaneously. **(a)** recorded with analog mode (showing bright areas of one image become dark areas of the other image) and **(b)** photon counting (showing almost no crosstalk between areas).

### Supplementary Note 1: PURELIGHT correction

In order to make use of all of the events measured during a TCSPC experiment, it is necessary to take into account all of the distortions that arise due to the pile-up effect. We can group these distortions into two categories: pile-up losses and time walk. In a threshold-based system where events are tagged when the analog signal goes over a set value, we can clearly see both distortions at play when looking at the uncorrected decay obtained from a high-count-rate simulation (Supplementary Figure SN1.1).

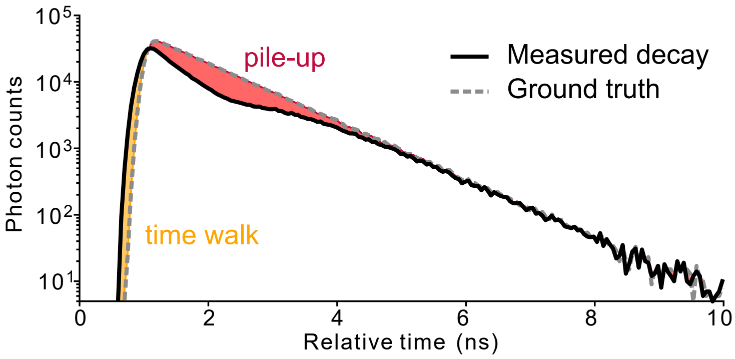

Supplementary Figure SN1.1: TCSPC simulation decay showing pile-up and time walk effects.

First, there is a clear pile-up loss effect distorting the exponential decay region (after 1.5 ns). Since this distortion consists in an arrival time-dependent loss of photon events, it affects both the lifetime estimation and the intensity of the measurement. The origin of the pile-up effect is the fact that, since the photon event is assigned every time the signal crosses a threshold, it is not possible to assign a second photon unless the signal has returned below the threshold level. Effectively, this defines a *blind time* for photon detection. In the ideal case of square pulses, the blind time is equal to the time that the signal spends above threshold, which we define here as *pulse width*.

For real pulses the rising and falling edges must also be considered (Supplementary Figure SN1.2). Due to the finite slope of the pulses, when two pulses are sufficiently close in time, the signal cannot return below baseline due to the summed contribution of the falling edge of the first pulse and the rising edge of the second. In the case in which the two pulses are actually present, the electronics detects one single pulse with a long pulse width. However, even if the second photon is not present, it is necessary to consider that if it would arrive, the electronics would not be able to detect it. For this reason, it is necessary to increase the blind time of each detected event by a fixed factor, here named "static blind offset" (SBO).

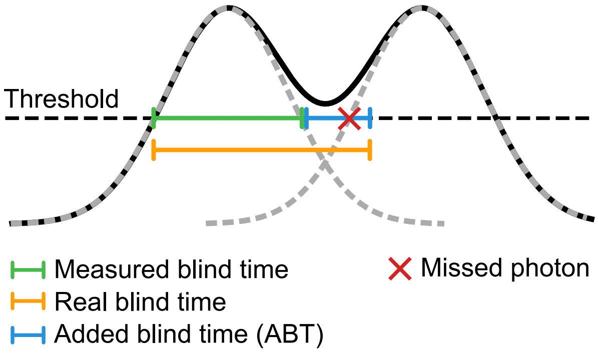

Supplementary Figure SN1.2: Origin of the ABT.

The second type of distortion, time walk, is evident when comparing the rising edge of the simulated detected decay in Supplementary Figure SN1.1 against the ground truth.

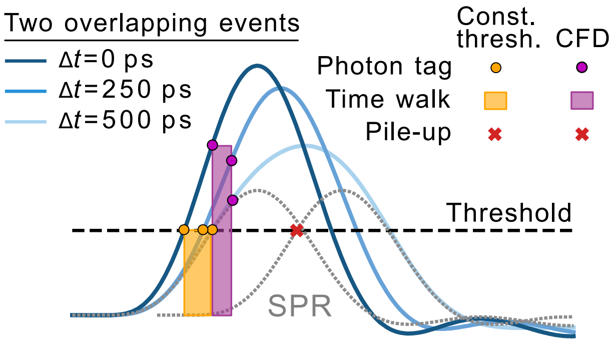

Supplementary Figure SN1.3: Time walk effect in constant threshold and constant fraction discriminator time tags.

The source of this effect is illustrated in Supplementary Figure SN1.3; the pile-up of multiple photon events that are very close in time affects the rise time of the analog signal leading to an early threshold crossing. In this case, the time shift is also associated to the failure to count the second photon event, i.e. to the photon losses. This occurs in all detector types, however the effect is particularly pronounced in detectors where the rising edge of the analog signal is relatively slow such as the SiPM. Additionally, for SiPMs there is a second source of time walk: the so-called "direct crosstalk" causes a certain percentage of detected photons to have a pulse amplitude that is a multiple of that of a single photon. This effect occurs at all count rates, generating distortions of the decays even at low light intensities.

#### PURELIGHT algorithm

The PURELIGHT correction combines three algorithms that together compensate for all the pile-up distortions described above:

1. **Added blind time (ABT)**: This is a pre-processing step that extends the blind time of each detected pulse beyond its pulse width. The minimum ABT is equal to the SBO, but often it must be extended further to account for secondary features of the pulse shape.

2. **Blind time compensation (BTC)**: This is an algorithm that corrects the photon loss due to the blind time by scaling the contribution of each photon event to the decay histogram by the fraction of time the electronics was able to detect photons in the temporal bin to which the photon was assigned.

3. **Time walk compensation (TWC):** This is an algorithm to correct the time walk effect. It estimates the shift for each photon event based on the additional information available in the XTTTR format and shifts the event back accordingly.

##### Added blind time (ABT)

The first step to apply the PURELIGHT correction is the added blind time algorithm. The ABT iterates over all the detected events $p \in P$ and sets the blind time ($BT_p$) equal to the sum of the pulse width ($PW_p$) and the predefined ABT value:

$$BT_p = PW_p + ABT \quad \forall\, p \in P. \tag{1}$$

If a photon event $p$ is within the blind time of a previous event $(p-1)$, it is removed, and the blind time of the $(p-1)$ event is adjusted to encompass the union of both events

$$BT_{p-1} = \max\left(BT_{p-1}, \Delta T_p + BT_p\right) \tag{2}$$

where $\Delta T_p$ is the time interval between the absolute times of the two photon events.

**Blind Time Compensation (BTC)**

Once the blind time for each photon event is set, the blind time compensation weights can be computed. These compensate the distortion generated by the blind time by scaling the detected events $p \in P$ by the ratio $\frac{N_{total}}{N_{active}(t_p)}$, where $N_{total}$ is total number of excitation cycles and $N_{active}(t_p)$ is the number of excitation cycles where the system was not blind for the relative time $t_p$ that corresponds to a detected event $p$. In practice the weights are computed as follows: First, compute the number of excitation cycles where the relative time $t$ was within a blind time period ($N_{blind}(t)$), with $t_{las}$ the excitation (laser) period:

**Algorithm 1** Computation of the blind cycles

```
N_blind(t) ← 0   ∀t ∈ [0, t_las)
for p ∈ P do
    for t_i ∈ [t_p, t_p + BT_p] do
        t'_i ← t_i  mod t_las
        N_blind(t'_i) ← N_blind(t'_i) + 1
    end for
end for
```

Then, compute the weight for each relative time $t \in [0, t_{las})$ as

$$w(t) = \frac{N_{total}}{N_{active}(t)} = \frac{N_{total}}{N_{total} - N_{blind}(t)}. \tag{3}$$

The computed weights will later be used to assemble the lifetime decay. Instead of adding a discrete count for each photon p in the histogram, the weight $w(t)$ is added. This adjusts the contribution of each photon to compensate for the probability of being able to detect a photon with that relative time.

**Time walk compensation (TWC)**

In the absence of pulse pile-up (i.e., when all photon events are non-overlapping in time) the ideal time tagging strategy would be to implement a constant fraction discriminator (CFD). This way of time tagging compensates for the variations in pulse amplitude (e.g. in PMT detectors) by tagging the time not at a constant threshold, but at a set fraction of the pulse height. However, in the presence of pile-up, this approach is incompatible with the BTC algorithm. For an analog pulse that incorporates contributions of several photon events, assigning the time tag using a CFD would result in tagging time that is a linear combination of the real times of arrival for the photons contained within that pulse. This would contradict the base assumption of the BTC method that once the analog signal is over the set threshold, the electronics are blinded, and no photon events will have a time tag within this period. To estimate the correct arrival time $t'_p$ for any photon event p we propose a general nonlinear function approach $f_{TWC}(p)$:

$$t'_p = t_p + f_{TWC}(p) \tag{4}$$

Where the function $f_{TWC}(p)$ uses the XTTTR information (e.g., pulse height, pulse width, slope at threshold crossing, area over threshold, etc.) of photon p to compute a correction factor for its arrival time. The function $f_{TWC}(p)$ can have different implementations, but all of them will make use of precomputed correction parameters which depend on the pulse shape characteristics of the detector. To obtain the correction constant, an instrument calibration procedure must be performed, as described in the corresponding section. We found that a simple yet effective implementation of the correction function, $f_{TWC,A_p}(p)$, consists of applying a piecewise linear shift $\Delta t_p(A_p)$ based solely on the area above threshold of the rising edge $A_p$:

$$\Delta t_p(A_p) = \begin{cases} \dfrac{\Delta t_{i+1} - \Delta t_i}{A_{i+1} - A_i}(A_p - A_i), & A_i \leq A_p \leq A_{i+1}, \\ \text{(defined analogously for other intervals)} \end{cases} \tag{5}$$

where $A_i$ and $\Delta t_i$ $\forall i$ are detector dependent and can be defined during calibration.

More advanced implementations of this function could also prove relevant in the future, including non-linear parameters and multiple pulse characteristics (e.g. pulse width, slope at threshold or any number of voltage values at arbitrary time intervals after threshold crossing). One such example could be a pre-trained neural network that combines all the information contained in the XTTTR format.

##### Combined correction

Finally, the lifetime decay histogram D is assembled by adding the weight of each photon event to the time bin corresponding to the corrected arrival time $t'_p$.

**Algorithm 2** Computation of the PURELIGHT decay

```
D(t) ← 0   ∀t
for p ∈ P do
    D(t'_p) ← D(t'_p) + w(t'_p)
end for
```

An example of the impact of each step of the correction on measured monoexponential data is shown in Supplementary Figure SN1.6.

#### Calibration of correction parameters

To determine the parameters for the PURELIGHT correction, we used the same fluorescein solutions and $KH_2PO_4$ crystals described in the methods to obtain a range of monoexponential lifetimes. For each solution we acquired a low count rate measurement (<5% of excitation rate) to obtain a reference decay and several high count rate measurements up to >100% of excitation rate or up to detector saturation. Then the following calibration procedure was performed:

##### Detection threshold, ABT, and $A_p$ width ($t_{A_p}$)

The detection threshold sets the voltage that must be crossed for a detection event to be recorded. Setting it too low can introduce fake photon counts from analog noise or low amplitude after-pulses, while setting it too high can lead to missed counts and increased time walk. Using the SPR of the detector (measured with a fast oscilloscope or by capturing data traces directly from the ADCs, Supplementary Figure SN1.4) one can set the detection threshold to 50% of the SPR peak amplitude. Once the detection threshold has been defined a value for the ABT can be obtained from the SPR by measuring the

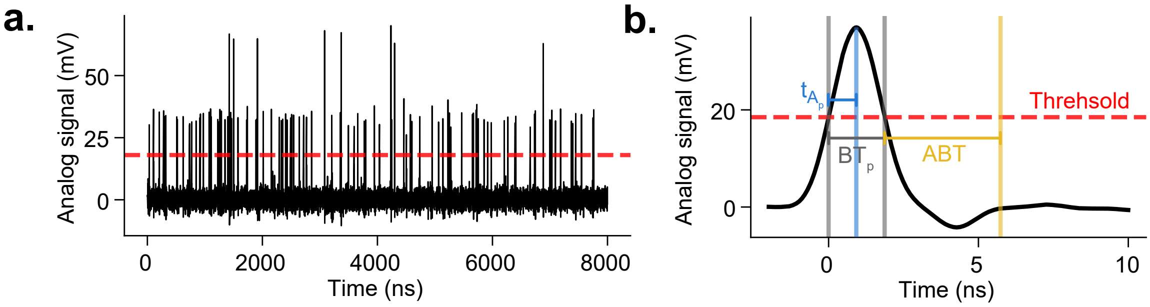

Supplementary Figure SN1.4: **Detection threshold and initial ABT estimation a.** ADC data trace (∼8 µs) showing single and double photon pulses and the event detection threshold. **b.** Custom SiPM detector SPR with the detection threshold set to 50% of the peak amplitude. The ABT is set to extend the blind time up to the point where the analog signal stabilizes after the pulse (∼4 ns in this case). The width of the integration time for $A_p$ ($t_{A_p}$) was set such that it would include the rising edge of the SPR from the threshold crossing up to the peak voltage.

time elapsed between the downward threshold crossing and the voltage signal stabilizing near 0 (Supplementary Figure SN1.4b).

##### $f_{TWC}(A_p)$ parameters

For the $f_{TWC}(A_p)$ it is necessary to define the width of the integration area used to measure $A_p$. This parameter needs to be set before data acquisition as $A_p$ is integrated directly from the digitized data within the FPGA. We found that setting $A_p$ to include the rising edge of the SPR from the threshold crossing up to the peak voltage (Supplementary Figure SN1.4b) provided a noise-robust and effective proxy for time walk compensation.

Next, the time-walk compensation parameters are estimated from a high-count-rate SHG measurement. Because all photon events in an SHG measurement should arrive at the same relative time (defined by the IRF), time walk can be measured directly as the shift in mean relative time as a function of $A_p$. A 2D histogram of relative time ($t_p$) against the area under the rising edge of each event ($A_p$) reveals clusters corresponding to different numbers of overlapping photons (Supplementary Figure SN1.5), and we define the time-walk compensation function $f_{TWC}(A_p)$ as a piecewise linear fit connecting these clusters. This yields the optimal correction for perfectly overlapping events, but overcompensates the timing drift of partially overlapping events, which dominate most fluorescence imaging scenarios. We therefore tune the compensation further by applying it to high-count-rate fluorescein measurements and inspecting the rising edge of the decay for distortion. Scaling the time-walk compensation by a factor of 0.65 gave the best rising-edge correction across all tested lifetimes and detectors.

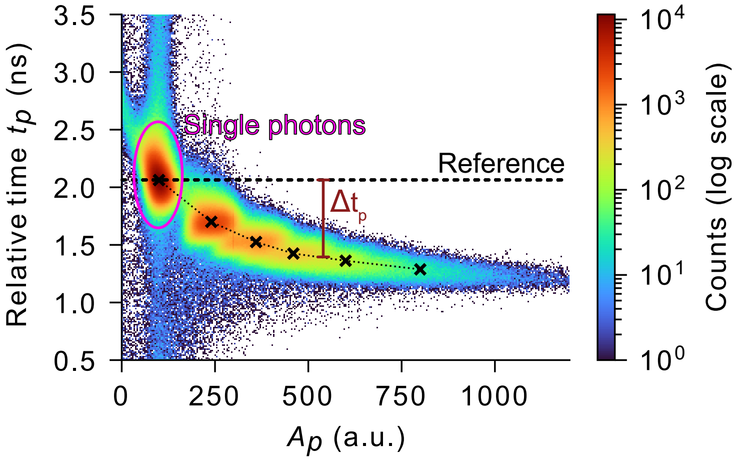

Supplementary Figure SN1.5: **Time walk compensation parameters.** A piecewise linear function maps the time walk as the $A_p$ increases. The first point of the function is taken as the reference to compute $\Delta t_p$ values.

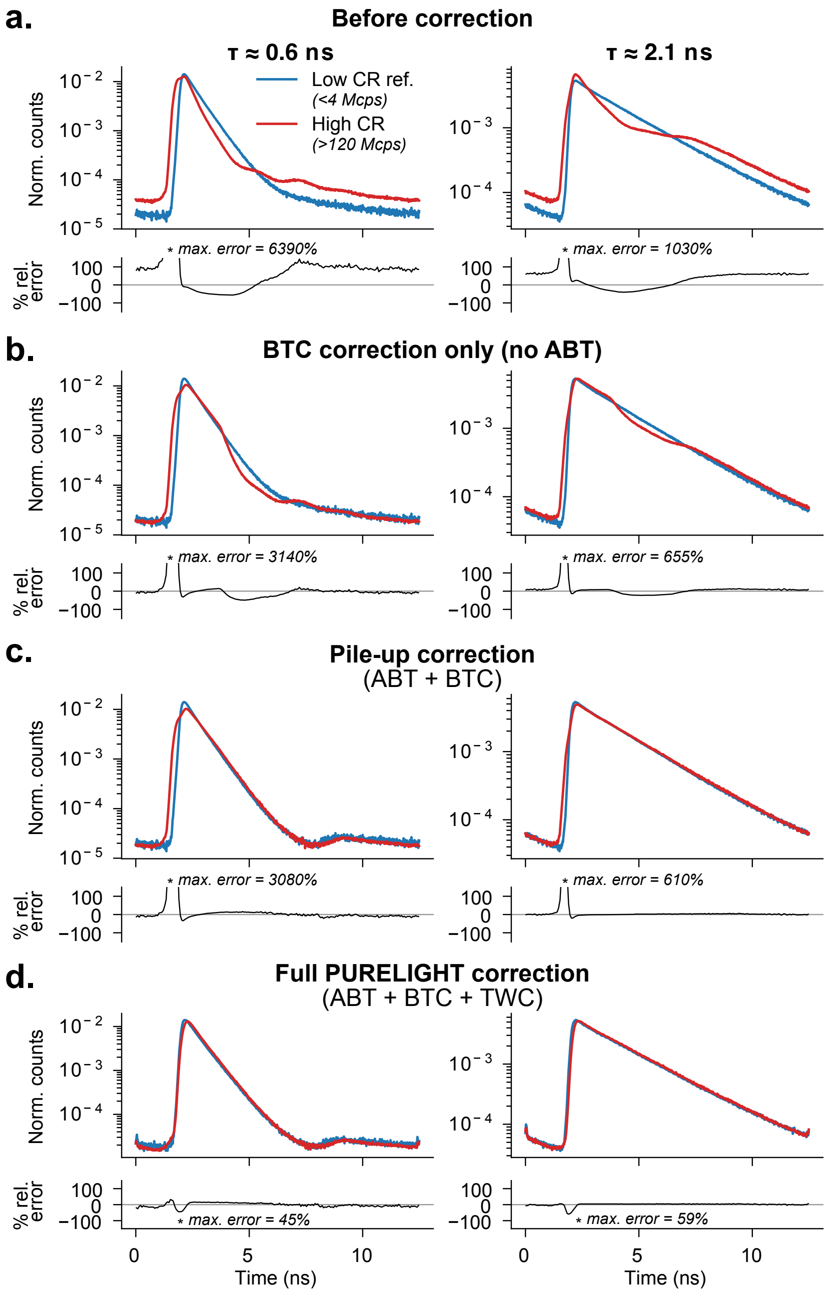

Supplementary Figure SN1.6: **Steps of the PURELIGHT correction on SiPM data. a.** Uncorrected TCSPC decays of two fluorescent solutions collected with the SiPM detector at low (blue) and high (red) count rates. The relative error plot (black) shows the difference between the area-normalized low and high count rate decays as a percentage of the low count rate values. **b.** Applying the blind time compensation (BTC) without the added blind time (ABT) only corrects part of the pile-up losses. **c.** Applying the BTC with the appropriate ABT fully corrects the pile-up losses resulting in matching decay slopes at low and high count rates. Even though the photon losses have been compensated by the BTC, the rising edge of the two measurements still shows large discrepancies due to time walk. **d.** Applying the time walk compensation (TWC) greatly improves the alignment of the rising edge of the low and high count rate measurements and concludes the full PURELIGHT correction.

#### Supplementary Figures

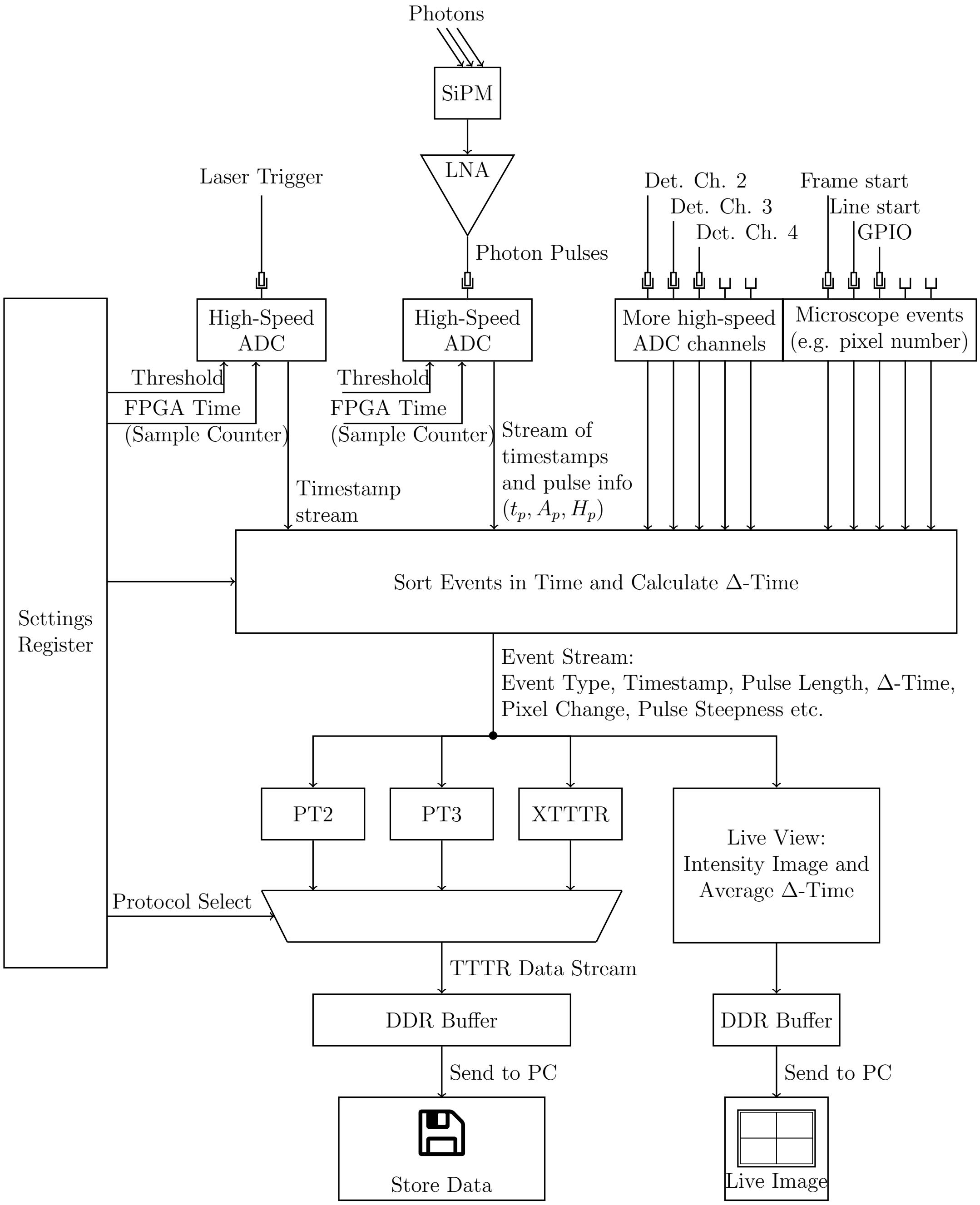

Supplementary Figure 1: **TCSPC implementation on the FPGA.**

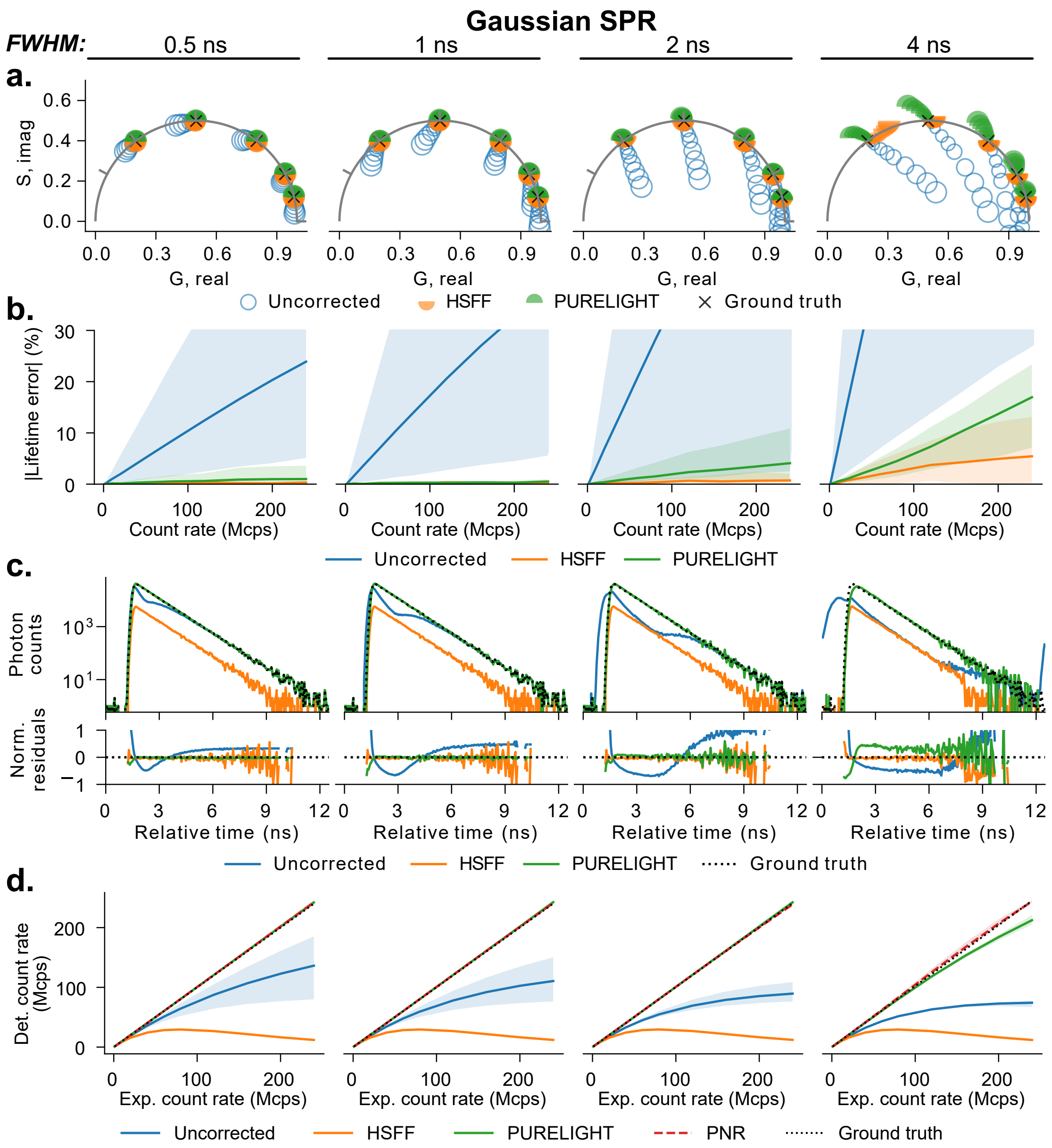

Supplementary Figure 2: **Validation of the PURELIGHT correction across different single photon detector response widths.** We simulated the detection of XTTTR data for a series of monoexponential lifetimes (0.25-4 ns + IRF) and count rates (1-240 Mcps; 1.25-300% of the excitation rate) with a Gaussian single photon response with FWHM of 0.5, 1, 2, and 4 ns. We then applied the HSFF and PURELIGHT corrections and compared to the ground truth and uncorrected data. **a)** Impact of count rate (illustrated as marker size) on the uncorrected and corrected phasor data for all simulated lifetimes. **b)** Relative lifetime error vs. count rate. The lines show the mean across all lifetimes, while the shaded areas show the range of errors across all lifetimes. **c)** Examples of the detected decays (top) and normalized residuals compared to the ground truth (bottom) for a 1 ns lifetime at 160 Mcps (200% of the excitation rate), illustrating the impact of the different corrections. **d)** Count rate linearity for the different corrections, showing the mean (solid/dashed line) and range (shaded area) across all lifetimes.

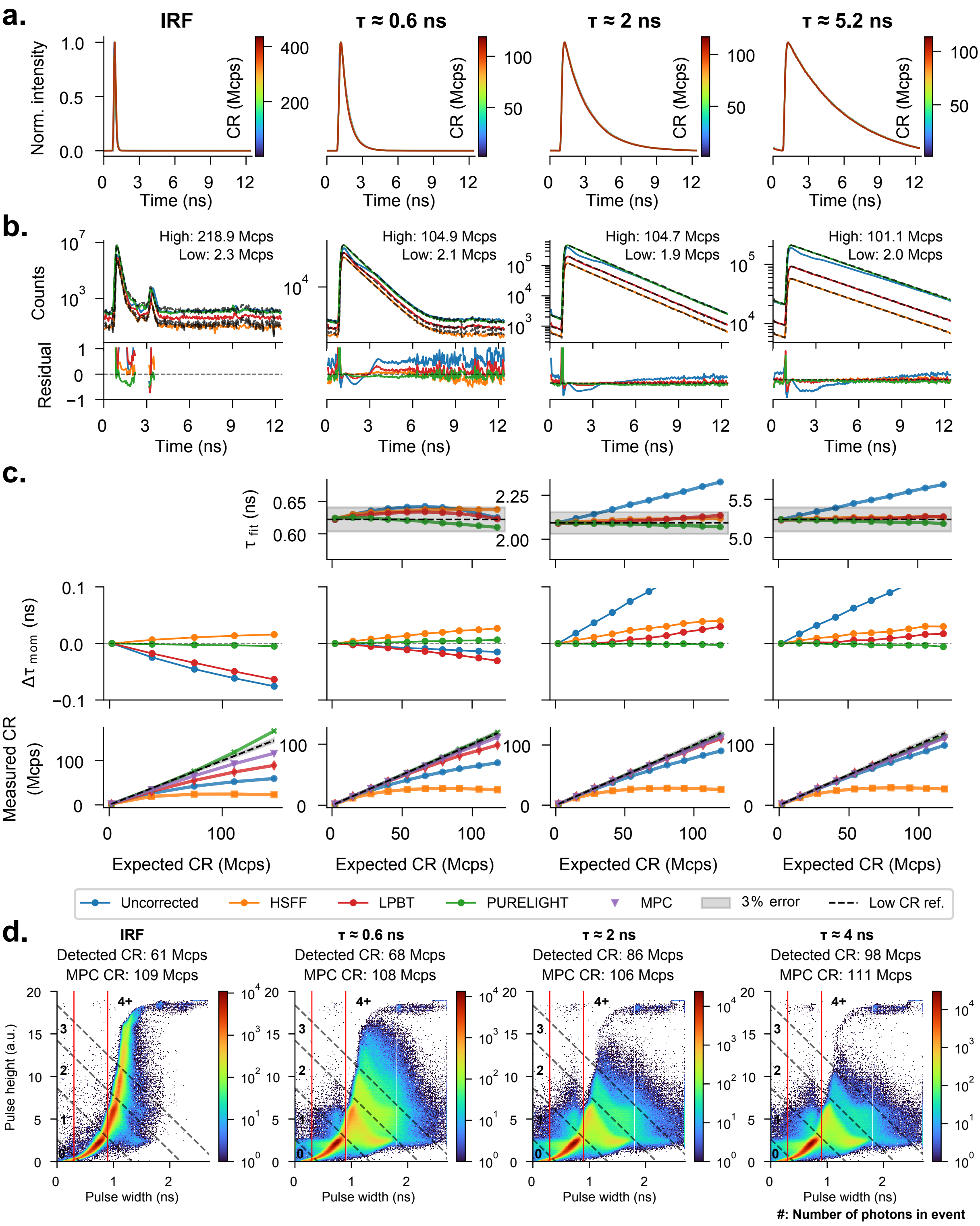

Supplementary Figure 3: **Hybrid Photodetector (HPD, PicoQuant PMA-40mod) PURELIGHT validation.** Continues on next page.

Supplementary Figure 3: **a.** PURELIGHT-corrected temporal decays of second harmonics generation (SHG, left) and fluorescent solutions with different lifetimes. Decays at multiple excitation powers are shown, normalized by the maximum and color-coded based on the estimated incident count rate. Fully overlapping curves indicate no variation in the measured fluorescence lifetime with increasing photon flux. **b.** Comparison between the different correction methods and the low count rate reference decay. For each correction method the reference was scaled to match the maximum counts and the residuals (bottom) were computed for time bins where the low count rate reference was above the dark counts limit. **c.** Fitted lifetime (top), method-of-moments lifetime drift (middle) and measured count rate (bottom) as a function of expected count rate for the different correction methods. The expected count rate was extrapolated based on a quadratic power dependency (linear regression with respect to the square of the power) estimated using the first four PURELIGHT corrected measurements. **d.** Pulse width vs. height heatmap of detection events for high count rate measurements with samples of different monoexponential lifetimes. The vertical red lines show the range of pulse widths that is considered a single photon detection event for the HSFF correction. The dashed black lines show the manually defined MPC classification boundaries.

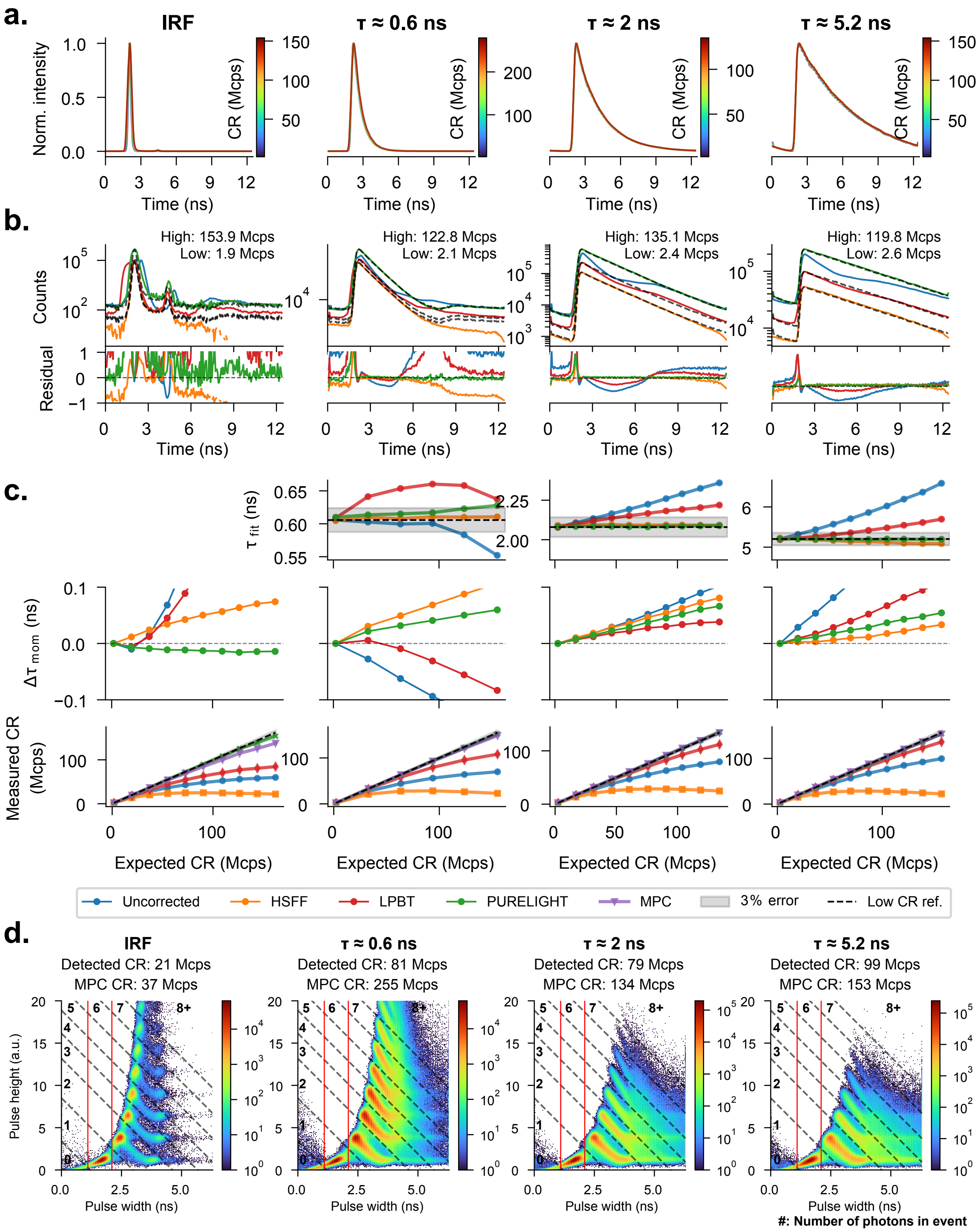

Supplementary Figure 4: **Custom silicon photomultiplier (SiPM) PURELIGHT validation. a-d.** Same as Supplementary Figure 3.

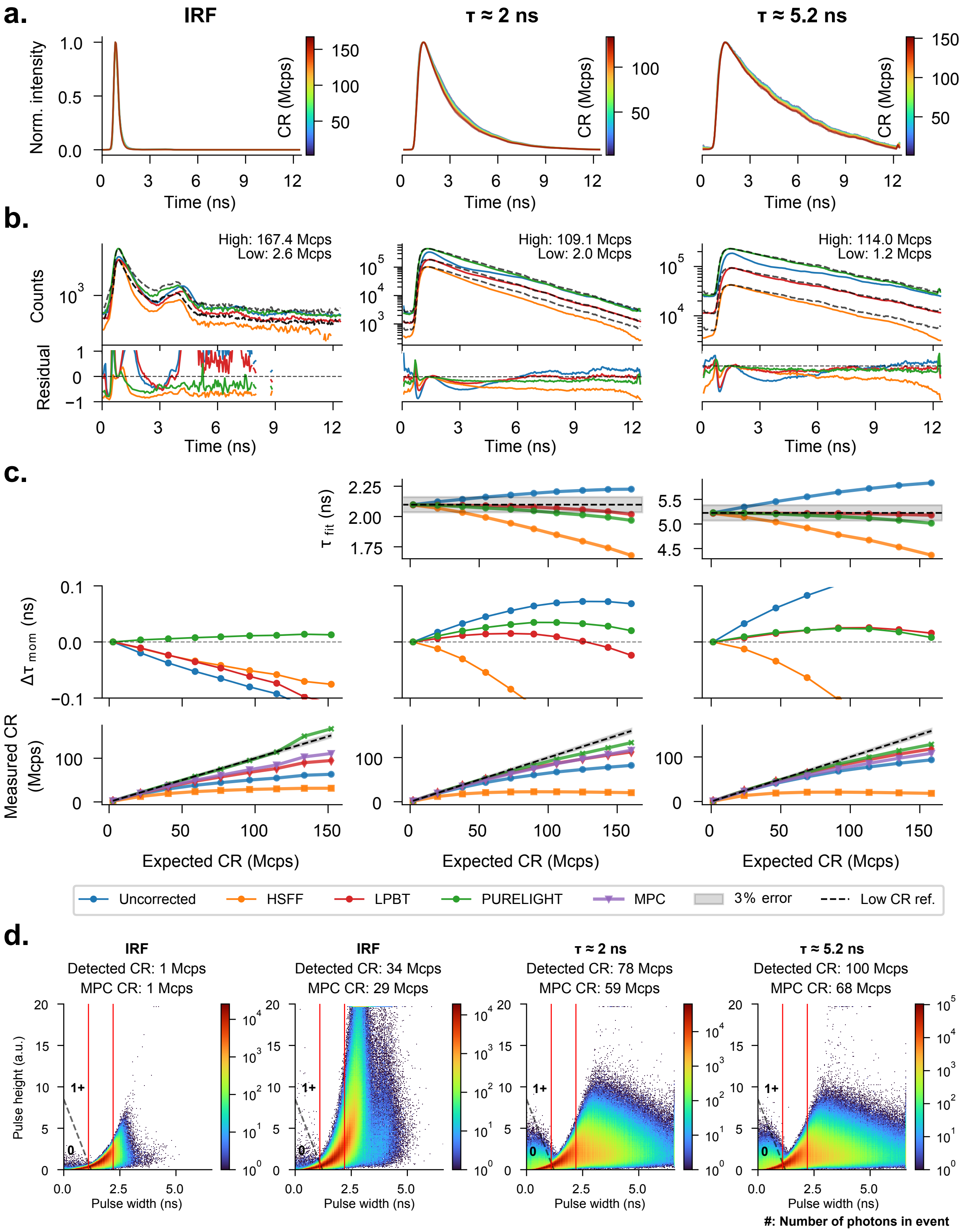

Supplementary Figure 5: **Photomultiplier Tube (PMT, Hamamatsu H10770PA-40) PURELIGHT validation. a-d.** Same as Supplementary Figure 3.

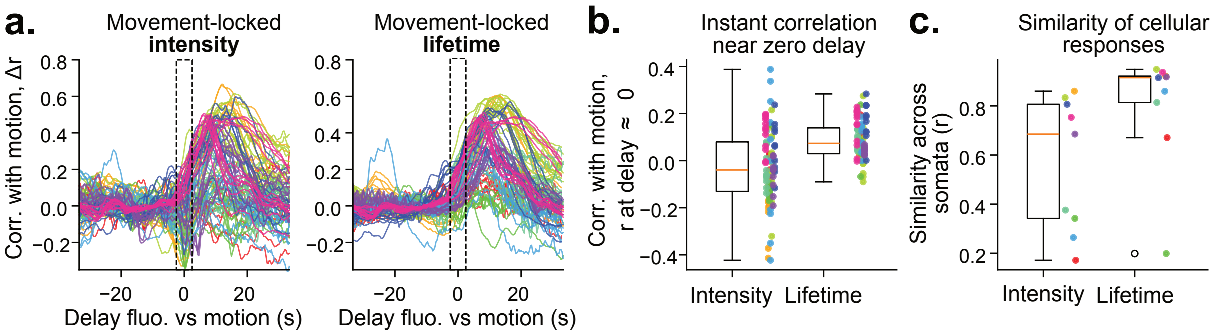

Supplementary Figure 6: **Lifetime-based astrocytic calcium measurements reduce motion-artifacts and provide more consistent responses across somata**. **a)** For each astrocyte soma, its fluorescence trace was correlated with brain motion extracted from the frame-to-frame image displacement estimated during motion correction (93 somata from 9 recordings in 3 mice; 180 s per recording, 3 Hz). Each line is one soma, colored by recording; curves are plotted relative to their average at negative delays, where they are flat (intensity r = -0.04, lifetime r = -0.02). **b)** Correlation averaged over a 5 s window centered on zero delay (dashed box in panel a). Intensity scatters around zero with some traces showing a negative correlation to motion, likely an artifact of the focal plane shift (median r = -0.04, IQR -0.13 to +0.08; 41% positive). Lifetime is consistently positive (median r = +0.07, IQR +0.03 to +0.14; 88% positive), reflecting the onset of the physiological response rather than motion contamination. Each dot is one soma, colored by recording; boxes show median and interquartile range. p = 0.004, Wilcoxon signed-rank on per-recording means (n = 9). **c)** Similarity of the delay curves in panel a between somata of the same recording, quantified as the median pairwise correlation. Lifetime responses are highly consistent from cell to cell (median r = 0.92 across recordings) and more consistent than intensity (r = 0.69; p = 0.03, same test). One dot per recording.

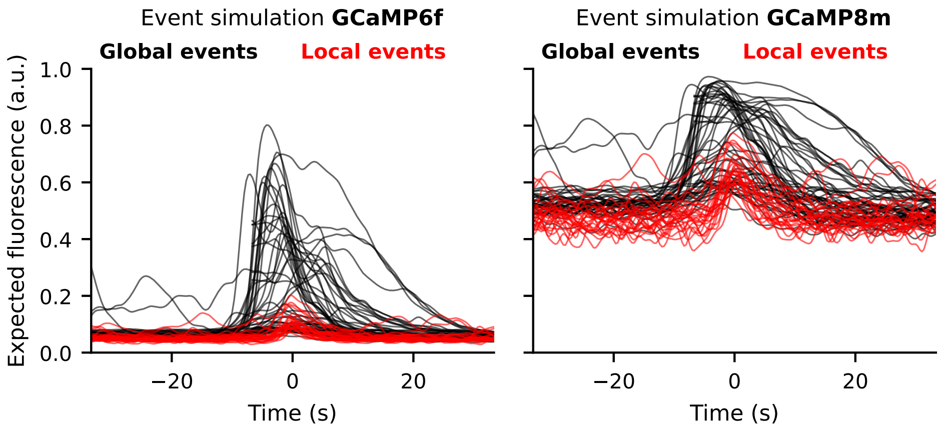

Supplementary Figure 7: **Simulated GCaMP6f and GCaMP8m intensity responses**. Expected fluorescence transients for large population events (global) versus cell-specific events (local) for standard intensiometric calcium indicators GCaMP6f and GCaMP8m. Events in absolute calcium concentration are taken from Figure 4f and converted to expected fluorescence using the Hill equation for the fluorescence $F = [\mathrm{Ca}^{2+}]^n/([\mathrm{Ca}^{2+}]^n + K_d^n)$, with Hill coefficient $n = 2.27$ and $K_d = 375$ nM for GCaMP6f, and $n = 1.92$ and $K_d = 108$ nM for GCaMP8m. The lower $K_d$ value of GCaMP8m results in two consequences: first, a higher baseline level, and second, a higher relative amplitude of local events compared to global events. Thus, GCaMP8m with its lower $K_d$ value improves the detection of local events in astrocytes.
